# Single-cell RNA sequencing of peripheral blood defines two immunological subtypes of Sjögren’s disease

**DOI:** 10.1101/2025.02.27.640483

**Authors:** Geoffrey Urbanski, Kimberly E. Taylor, Emily Flynn, Armita Norouzi, Catherine Chu, Brittany Davidson, Arnab Ghosh, Joanne Nititham, Ravi K. Patel, Annie W. Poon, Gabriela K. Fragiadakis, Alexis J. Combes, Walter L. Eckalbar, Lindsey A. Criswell, Chun Jimmie Ye, Caroline H. Shiboski

## Abstract

Sjögren’s disease (SjD) is a heterogeneous autoimmune disorder with significant clinical and molecular diversity. While anti-SSA antibodies serve as a hallmark serological biomarker, nearly half of patients lack them, raising fundamental questions about distinct pathogenic mechanisms. To address this, we performed single-cell RNA sequencing with surface protein profiling on 1.5 million PBMCs from 333 participants in the Sjögren’s International Collaborative Clinical Alliance cohort, comparing those with and without SjD and stratifying them by anti-SSA status. Our analysis identified two immunologically distinct subtypes of SjD, with SSA-positive patients exhibiting a dominant and persistent IFN-I signature, profoundly impacting immune cell maturation. Notably, transitional B cells were disproportionately affected, displaying signs of premature maturation, reduced BCR diversity, and shorter CDR3 regions, reinforced interactions with proinflammatory cells, thereby fostering an environment conducive to autoreactivity. By contrast, SSA-negative SjD participants lacked an upregulated IFN-I signature, challenging prevailing pathogenic models and suggesting alternative immune dysregulation pathways. These findings support a two-disease model of SjD and highlight transitional B cells as both a key biomarker and a potential therapeutic target. Targeting IFN-I signaling and transitional B cell maturation may represent a novel therapeutic avenue to modulate immune dysregulation and prevent autoreactivity in SjD.

## Introduction

Sjögren’s disease (SjD) is a multifaceted systemic autoimmune disease ^1^. Its primary hallmark is lymphocytic infiltration of the exocrine glands, leading to pronounced dryness of the mouth and eyes. However, SjD exhibits significant systemic heterogeneity, with 30–40% of patients developing extraglandular manifestations (EGM) affecting the joints, peripheral and central nervous system, lungs, skin, kidneys, muscles, and hematopoiesis ^1^. Critically, patients with SjD have a 7- to 19-fold increased risk of developing lymphoma compared to the general population ^2^. SjD diagnosis is guided by classification criteria ^3^, with anti-SSA (anti-SSA60kDa/Ro) antibodies serving as the hallmark serum biomarker for the disease. However, unlike other connective tissue diseases where autoantibodies are rarely absent, SjD is unique in that nearly half of patients lack detectable anti-SSA antibodies ^4,5^. This creates a dichotomy in SjD with SSA-positive and SSA-negative patients, which impacts the disease phenotype as the presence of anti-SSA antibodies has been strongly associated with multiple factors: younger age at diagnosis ^6^, higher frequency of EGM ^6,7^, and increased markers of B-cell activation, including cryoglobulinemia ^5,7^, complement consumption ^5^, rheumatoid factor presence ^7^, and hypergammaglobulinemia ^7^.

The precise pathogenesis of SjD remains poorly understood, with the main hypothesis focusing on exocrine gland epithelium, where immune cells accumulate, leading to the formation of lymphoepithelial lesions ^8,9^. In this hypothesis, the interferon (IFN) signaling pathway plays a critical role: After stimulation, Toll-like receptors (TLR) 3, 7, and 9 ^10^ induce type I IFN (IFN-I) response by activating IFN-stimulated genes (ISG), the Janus kinase/signal transducer and activator of the transcription 1 pathway, and the gene expression of many inflammatory molecules ^11^, including B-cell activating factor belonging to the TNF family (BAFF), a key cytokine for B cells in SjD ^12^. The IFN-I-induced proteins increase apoptosis and the release of self-antigens, stimulating antigen-presenting cells, which express MHC class II molecules ^13^. Genome-wide association studies support this, as most susceptibility loci were located in ISGs and class II HLA genes ^14,15^. However, despite this central role attributed to IFN in SjD pathogenesis, nearly one-third of SjD patients lack elevated blood IFN signature ^16^. Interestingly, the IFN signature is much more prevalent in patients who produce anti-SSA antibodies ^17,18^. However, the absence of circulating molecular markers in SSA-negative individuals goes against the current pathogenic hypothesis, since immune cells are expected to originate from blood before infiltrating exocrine glands. Better cellular and molecular characterization of circulating immune cells is critical for the diagnosis and stratification of SjD and the development of course-reversing treatments ^19^.

Single-cell RNA sequencing (scRNA-seq) of Peripheral Blood Mononuclear Cells (PBMCs) provides a unique opportunity to investigate the transcriptional state of individual circulating immune cells. To date, PBMC scRNA-seq studies in SjD have been greatly constrained by sample sizes, ranging from 5 to 24 samples, limiting the investigation of disease heterogeneity ^20–26^. In this study, we aimed to investigate the PBMC landscape of SjD at single-cell resolution, leveraging the large biorepository from the Sjögren’s International Collaborative Clinical Alliance (SICCA). We sought to delineate disease-specific immune signatures by comparing SjD participants to non-SjD symptomatic controls and examining differences between SSA-positive and SSA-negative participants to uncover distinct pathogenic mechanisms underlying these two subtypes.

We generated a comprehensive multi-omics dataset, including gene expression profiles, 132 cell surface markers, and BCR/TCR sequencing from 1.5 million PBMCs across 333 unique patient samples from the SICCA cohort ^3^, using Single Cell 5′ transcriptomic sequencing. Our analyses revealed virtually no significant differences in gene expression between SSA-negative SjD (SSA-SjD) and SjD-negative (SjD-) participants. By contrast, analysis of the multimodal data revealed substantial differences in SSA+SjD samples compared to both SSA-SjD and SjD- individuals. These differences were characterized by increased expression of IFN-I ISGs in each category of cells (T/NK, monocytes, B), and had a significant effect on B cell maturation. Targeted analysis of B cells revealed an expansion of immature circulating B cells, reduced B cell diversity, and shorter CDR3 heavy chains, fostering an environment conducive to autoreactivity. Joint analysis of molecular features also revealed an inverse correlation between B cell diversity and increase of type-I IFN response in SSA+SjD.

## Results

### Sample selection and group characteristics

We used multiplexed droplet-based scRNA-Seq (Figure 1A) and CITE-Seq to profile 1,537,678 PBMCs from 423 samples from the University of California, San Francisco (UCSF) SICCA site (Flow chart in Supplemental Figure 1), including 90 follow-up samples. The 333 baseline participants consisted of 101 SSA+SjD, 47 SSA-SjD, and 185 SjD- individuals. Overall, general demographic and phenotypic characteristics did not differ significantly (Table 1). Expectedly, SSA+SjD participants exhibited more extra-glandular manifestations, a higher frequency of anti-SSB antibodies, and greater B-lymphocyte signs of activity (BLA) compared to SSA- SjD participants.

**Figure 1.**
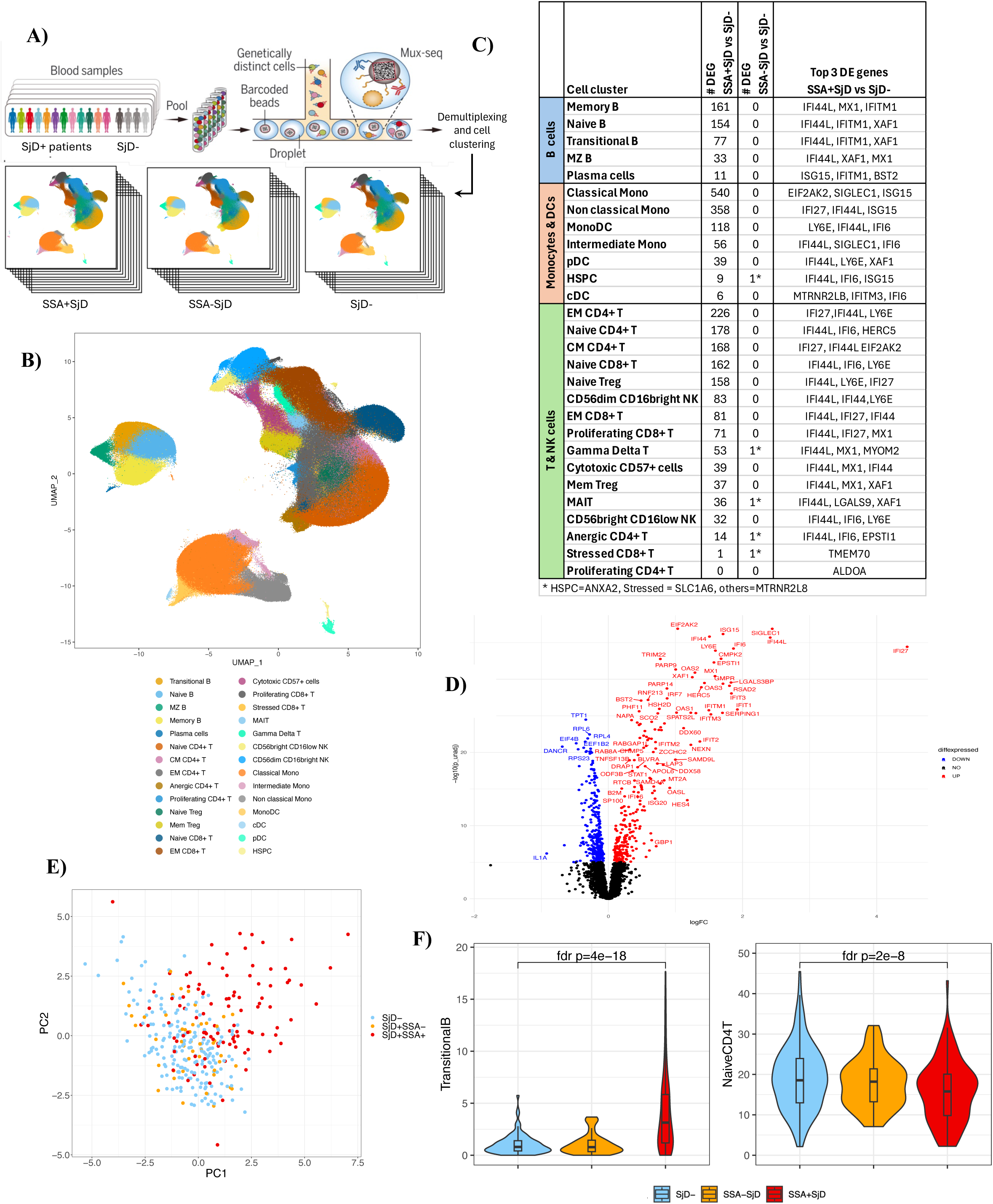
Differential expression and abundance are dominated by differences seen only in SSA+SjD. A) Overview of experiment: patient PBMCs are pooled, cells barcoded and sequenced, then patients identified via genetic demultiplexing and cell types identified via clustering gene expression; B) UMAP of Louvain clusters with cell type annotation; C) DEGs in SSA+SjD and SSA-SjD compared to SjD- in 28 cell clusters. Number of DEGs per analysis using EdgeR, and top three DEGs in SSA+SjD; D) Volcano plot of-log10(p-value) versus logFC of expression for SSA+SjD versus SjD- in classical monocytes; E) Principal components 1 and 2 from PCA of cell type abundances per sample; F) Differential abundances in top 2 cell types, Transitional B and Naïve CD4 T, with Dirichlet p-values.

**Table 1.**
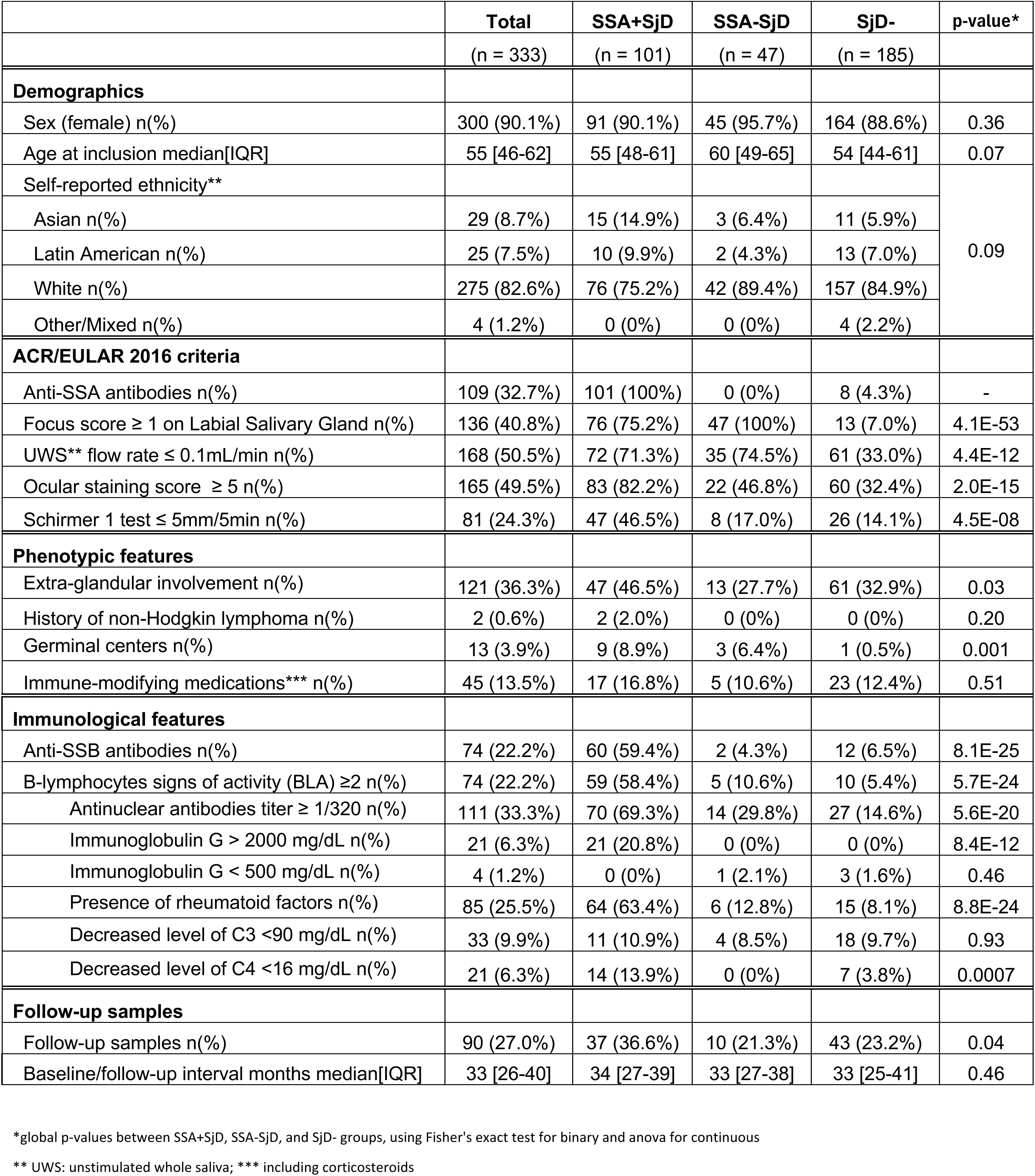
SICCA participant statistics.

### ISGs are upregulated in SSA+SjD but absent in SSA-SjD

Louvain clustering of all cells identified 28 PBMC clusters (Figure 1B, marker plot in Supplemental Figure 2). Surface protein expression from cells in three of the five batches was analyzed using 132 cell surface antigens (CITE-Seq) to validate finer annotations of RNA-based cell clusters.

To assess changes in the molecular states of each cell type, we performed differential gene expression (DE) analysis using EdgeR^27^ comparing SSA+SjD, SSA-SjD, and SjD- samples across the 28 cell types. We saw a striking difference across almost all cell clusters, with hundreds of differentially expressed genes (DEGs) in SSA+SjD and virtually no DEGs in SSA-SjD, compared to SjD- (Figure 1C). Most of the top DEGs for SSA+SjD were ISGs with the most frequently upregulated DEGs being *IFI44L*, *IFI27*, and *MX1*, which are markers of IFN-I response. Across cell types, there was a wide range of DEGs in SSA+SjD vs SjD-. Figure 1D illustrates this for classical monocytes, which have the highest number of DEGs. To account for differences in participant and cell numbers across groups, we compared log-fold changes (logFC) for SSA+SjD vs. SjD- and SSA-SjD vs. SjD-. This confirmed that the observed pattern reflected true effect size differences rather than variations in power (Supplemental Figure 3).

### Transitional B Cells are significantly expanded in SSA+SjD, driving differential cell abundance

We next examined differences in cell type abundance between the three groups. Principal components analysis (PCA) of the relative abundance of each cell type (Figure 1E) revealed a clear separation of the SSA+SjD group from the other two groups. The presence of anti-SSA antibodies was more strongly correlated with principal components PC1 and PC2 (r=0.46 and 0.36, respectively, p<5e-5) than SjD status (r=0.37 and 0.30, p<5e-5). There was no significant difference in PC1 and PC2 between SSA-SjD and SjD- participants. The cell type most strongly associated with PC1 was transitional B cells (r=0.71, p<5e-5). Differential abundance (DA) analysis via Dirichlet regression^28^ revealed significant differences in cell type proportions, again primarily in SSA+SjD compared to the two other groups (Supplemental Table 1). The most notable was a four-fold increase in the frequency of transitional B cells in SSA+SjD versus SjD- (SSA+SjD 4.0% versus SjD- 1.1%; p_fdr_=4e-18), followed by naive CD4+ T cells (15.8% versus 19.1%, p_fdr_=2e-8) (Figure 1F).

### Tensor decomposition analyses reveal correlation of molecular and cellular phenotypes

Single-cell transcriptome analysis offers a unique opportunity to simultaneously explore the dependencies between multiple cell composition and cell state phenotypes, which is challenging to capture through independent gene expression profiling by cell type. To assess this complexity, we applied single-cell tensor decomposition analysis (scITD)^29^, a computational method designed to extract cross-cell type gene expression programs, enabling detection of joint patterns of dysregulation. scITD produces an unsupervised summary of gene expression diversity across a finite number of factors; each factor has a loading for each gene-cell type combination and a donor score reflecting the degree of the factor present in each participant. Out of twelve factors, the factor explaining the most variance (Factor 1) stood out as the only one independently associated not only with SjD status but also key features of SjD, including the presence of anti-SSA antibodies, focal lymphocytic sialadenitis (with focus score (FS) ≥1), and the presence of anti-SSB antibodies (Figure 2A). We clustered genes highly associated with Factor 1 loadings (max absolute loading > 50 and gene-factor p-value < 5e-5) (Figure 2B). Out of the 42 genes, only 12 of the 38 upregulated genes were not identified as clear ISGs (*PLAC8, GIMAP4, SCO2, PARP14, APOBEC3A, MT2A, TNFSF10, EPSTI1, PLSCR1, STAT1, SAMD9L, HERC5, LGALS3BP*). We assessed correlation of the Factor 1 donor score (F1) with previously-published interferon modules ^30^: M1.2 (*IFI44, IFI44L, IFIT1, IFIT3, MX1*), M3.4 (*ZBP1, EIFAK2, IFIH1, PARP9, GBP4*), and M5.12 (*PSMB9, NCOA7, TAP1, ISG20, SP140*). These modules reflect the type I (M1.2 and M3.4) and type II (M3.4 and M5.12) IFN signatures. M1.2 served as a strong proxy for F1, showing a very high correlation (r=0.80). F1 also correlated highly with M3.4 (r=0.67) but moderately with M5.12 (r=0.43) (Figure 2C, 2D, Supplemental Figure 4A).

**Figure 2.**
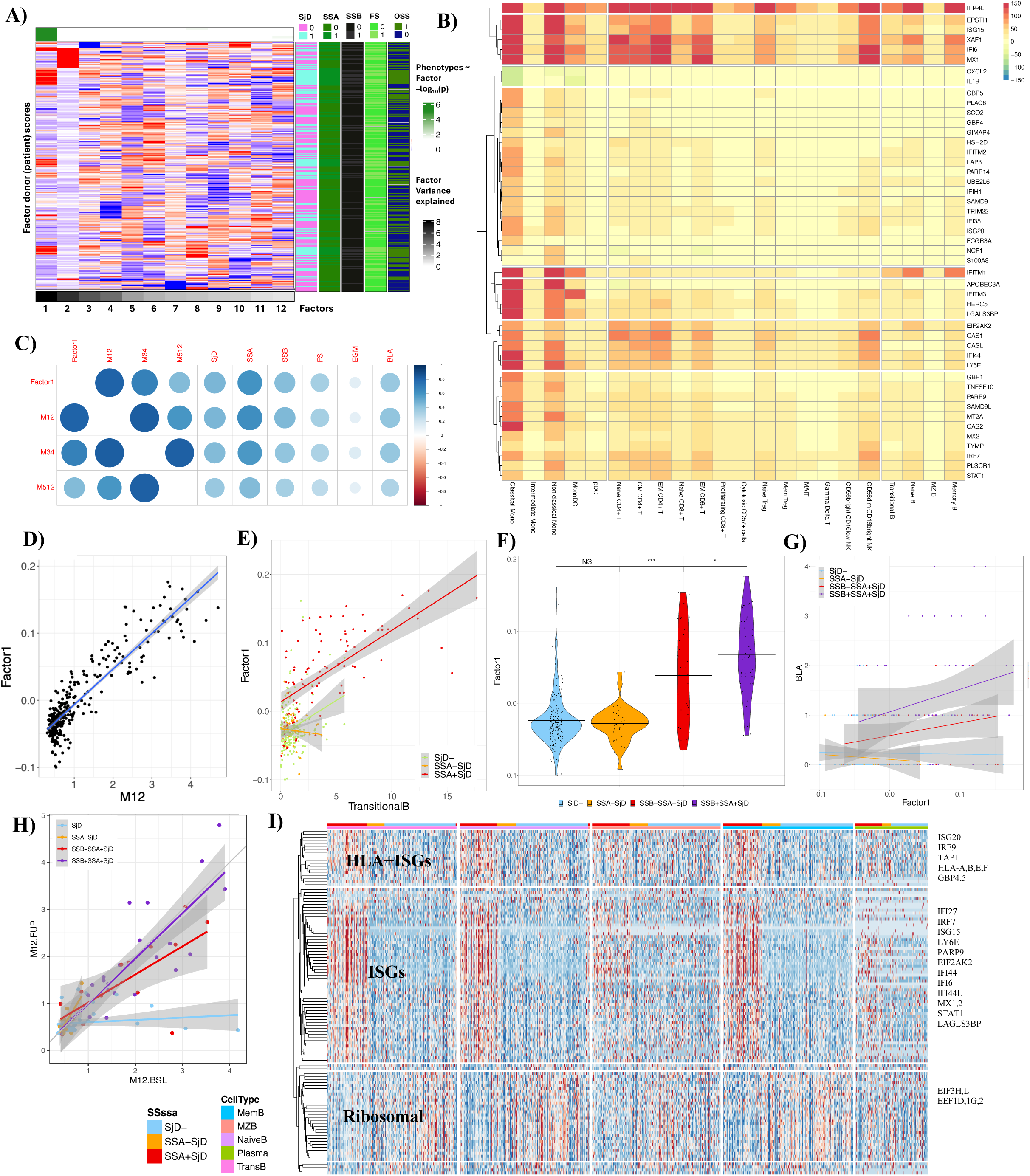
Single-cell interpretable tensor decomposition elucidates a cross-cell type pattern of ISG expression which is associated with SSB and B lymphocyte activity in the presence of SSA. A) scITD factors with donor (patient) scores and phenotype status in each row, top bar contains association (green significant) of factors with 5 phenotypes SjD, SSA, SSB, FS and OSS; B) Clustered top Factor 1 gene-cell type loadings (factor-gene p<5e-5, max absolute loading >50); C) Correlations of Factor 1 with IFN Chiche modules M1.2, M3.4, andM5.12, and phenotypes; D) Scatter plot of Factor 1 by M1.2 per sample; E) Scatter plot of Factor 1 and Transitional B abundance; F) Factor 1 by SjD/SSA/SSB groups; G) Scatter plot of Factor 1 and BLA, by SjD/SSA/SSB group; H) Scatter plot of M1.2 in baseline vs. follow-up samples, by SjD/SSA/SSB group; I) pseudo-bulk heatmap of clustered DEGs for B cell subtypes.

F1 was strongly associated with cellular composition (Supplemental Table 1), having the highest correlation with the proportion of transitional B cells (r=0.68, p=5e-40, Figure 2E) and CM CD4+ T cells (r=0.57, p=5e-26).

We next examined the correlation of the inferred factors with cellular composition and clinical features. F1 was more strongly correlated with the presence of anti-SSA antibodies (r=0.59, p<1e-14) than with SjD disease (r=0.45, p<1e-14) (Figure 2C). However, F1 score did not correlate with the presence of EGM (r=0.08, p=0.17), indicating a disconnect between clinical activity and gene expression. In addition to anti-SSA status, F1 was associated with the presence of anti-SSB and BLA (as defined in Methods) (Figure 2C). After stratifying SSA+SjD participants into two subgroups based on the presence (n=60) or the absence (n=41) of anti-SSB antibiodies, we observed a significantly higher F1 in the SSB+SSA+SjD subgroup compared to the SSB-SSA+SjD (Figure 2F). This aligns with more defined cases according to 2016 ACR/EULAR classification criteria score (Supplemental Figure 4B). We also confirmed a stronger correlation between the number of signs of BLA and F1 in the SSB+SSA+SjD group compared the two other SjD subgroups (SSB-SSA+SjD and SSA-SjD) (Figure 2G).

Multivariate modelling of F1 association with anti-SSA, anti-SSB antibodies, and BLA confirms that interactions of anti-SSA with anti-SSB antibodies or BLA are significant positive predictors of F1 (coefficients 0.041 and 0.029 with p-values 0.023 and 0.017 respectively) in addition to the main effect of anti-SSA antibodies (Supplemental Table 2). In joint models stratified by anti-SSA status, BLA and anti-SSB antibodies have significant associations with F1, but only in the presence of anti-SSA antibodies. These findings demonstrate a strong and proportional relationship between the activation intensity of IFN-I and the level of BLA, as evidenced by the production of anti-SSA and/or anti-SSB antibodies or by non-specific biological markers.

IFN-I signature remained stable over time, as indicated by comparing M1.2 in 90 participants between baseline and follow-up time points (median time interval of 33 [26-40] months) (Figure 2H). We also observed no significant differences in expression or abundances between these time points. This is consistent with the clinical evolution, as only 4/90 (4.4%) participants changed from SjD- to SjD. However, immunosuppressive/immunomodulatory drugs did impact ISG expression. We observed lower F1 in SSA+ participants who were under those treatments compared to those who were not (Supplemental Figure 4C).

### Heterogeneous IFN signature across immune cells suggests an impact in B cell differentiation

*IFI44L* appeared to have a homogeneous influence on Factor 1 across cell types (Figure 2B) but most other ISGs showed clear heterogeneity within major cell groups (such as monocytes, dendritic cells, NK cells, B cells, and T cells), and even within these specific immune cell types (Figure 2B), suggesting different effects of IFN on those subtypes. A heatmap of pseudo-bulk gene expression in B cells confirms this higher level of granularity (Figure 2I), highlighting three distinct gene modules (pathway analysis in Supplemental Figure 4D-E) differentially expressed in SSA+SjD samples compared to other groups. In B cells, we confirmed no significant differences in gene expression between SSA-SjD and SjD- specimens. The primary upregulated module was predominantly composed of IFN-I-related ISGs, while the second module included a broader set of ISGs, encompassing both IFN-I genes *(ISG20, IRF9*) and type II IFN response genes (*GBP4, GBP5, TAP1*). Additionally, this module featured antigen presentation genes, including both classical (*HLA-A, HLA-B*) and non-classical genes (*HLA-E, HLA-F*) MHC-I genes. Interestingly, MHC-I gene dysregulation varied across the five B cell subsets, and was also present in transitional B cells, which are supposed to have a very low antigen-presenting activity in normal conditions.

### Fine-grained clustering reveals immature B cell enrichment in SSA+SjD, driven by IFN-I stimulation

To further investigate the heterogeneity within major circulating immune cell types, we reclustered and deeply analyzed monocytes and dendritic (MD) cells, T and NK cells, and B cells separately.

Unsupervised Louvain-based cell clustering identified 27 different populations across MD cells (Supplemental Figure 5A). Four monocyte populations were significantly enriched in SSA+SjD specimens, exhibiting high ISG expression (*ISG15, IFI6, IFI44L, MX1, MX2*) compared to all other cell clusters: ISG+ AP1low classical monocytes (4.1-fold increase compared to SjD-, p_fdr_=6e-52)), ISG+ AP1 high classical monocytes (2.9-fold increase compared to SjD-, p_fdr_=9e-20), HLA-DR-high intermediate monocytes, and ISG+ non-classical monocytes (3.3-fold increase compared to SjD-, p_fdr_=9e-27) (Supplemental Figure 5B and Supplemental Table 3). Stimulation by IFN did not specifically favor any monocyte differentiation pattern. In addition, apart from the ISG-enriched monocyte populations, most other monocyte subsets were decreased in SSA+SjD samples, regardless of their polarization as M1 (CD163-, pro-inflammatory) or M2 (CD163+, anti-inflammatory) (Supplemental Figure 5C, Supplemental Table 3). In k-Nearest Neighbors (kNN) reclustering of MD cells using MiloR ^31^, which identifies finer-grained cell “neighborhoods” defined by expression similarity, no neighborhoods appeared to differ between SSA-SjD and SjD- groups, while neighbors enriched in SSA+SjD showed higher IFN expression (Supplemental Figure 5D-F).

Reclustered T/NK cells spanned 40 distinct populations (Supplemental Figure 6A). Similar to monocytes, a few clusters exhibited intense ISG expression and were abundant in SSA+SjD participants: ISG+ CD56bright CD16- NK cells, ISG+ GZMK+ effector memory CD8+ T cells upregulating *IFIT1, IFIT2, IFIT3, ISG15, and OASL*, and particularly ISG+ naive CD4+ T cells (3.2-fold increase in abundance compared to SjD-) upregulating *IFI44L, IFI6*, and *ISG15* (Supplemental Figure 6B, Supplemental Table 4). Neighborhood analysis highlighted that SSA+SjD samples exhibited all significant differences compared to SjD-, while SSA-SjD specimens showed no distinction from SjD- (Supplemental Figure 6C-D). However, it also revealed no specific pattern of neighborhood imbalance beyond increased maturation in naïve T cells with higher *KLF2* expression, greater activation of T cells marked by *ICOS*, and enhanced terminal differentiation of effector memory cells but neighbors enriched in SSA+SjD showed higher IFN expression (Supplemental Figure 6D-E). In addition, scRNA-seq TCR (VDJ) analysis revealed few differences across groups except for a shorter length of the beta chain exclusively in SSB+SSA+SjD samples (Supplemental Figure 6F).

Unsupervised B cell clustering identified 23 populations, revealing significant diversity within transitional B cells, which were defined by canonical markers (IGHD, CD24, CD38) and the expression of the pre-BCR component *VPREB3* (Figure 3A-B). Activated ISG+ Kappa transitional B cells and ISG+ Kappa naive B cells formed unique clusters (Figure 3B) and were much more abundant in SSA+SjD (Figure 3C, Supplemental Table 5). We identified distinct subtypes of transitional B cells, including follicular pathway-T2/T3 transitional (late transitional) cells, as well as Marginal Zone Precursors/T2 transitional (MZP-T2) cells, which progress into marginal zone precursors (MZP) (extra-follicular pathway). Similar to MD and T/NK cells, neighborhood analysis revealed enriched neighborhoods only in SSA+SjD participants (Figure 3D-E). This enrichment was primarily found in early forms of B cells, with a notable concentration among transitional B cells (Figure 3D). Additionally, across neighborhoods, higher SSA+SjD enrichment was strongly and positively correlated with M1.2 IFN score (Figure 3F).

**Figure 3.**
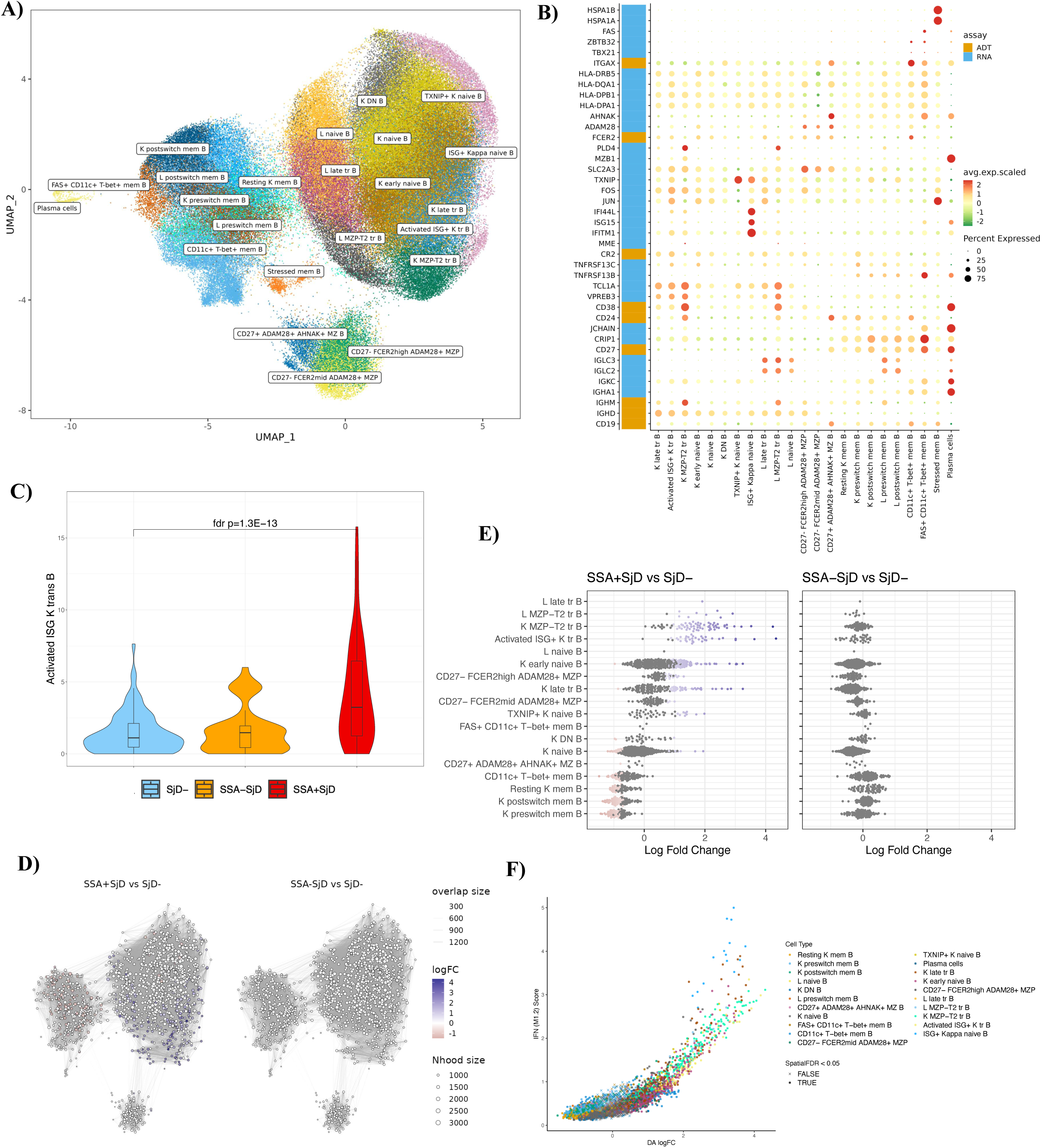
Fined-grained cell annotation and neighborhood analyses identify high-ISG subpopulations in B cells. A) UMAP of refined clusters within B cells; B) dotplot of marker genes for annotation of reclustered B cells; C) abundance of Activated ISG+ K transitional B cells by SjD/SSA group; D) Milo neighborhood differentials between SSA+SjD and SSA-SjD versus SjD- in B cells; E) Neighborhoods mapped onto the cell clusters with the highest percentage (if >30%) of cells, with logFC of differential abundance, purple=significantly higher than SjD-, pink=significantly lower than SjD-; E) Factor 1 by DA logFC of Milo neighborhoods.

### Anti-SSA antibodies are linked to immature B cell expansion, reduced diversity, and restricted antigen recognition

Single-cell BCR (VDJ) data showed significantly higher percentage of IgM isotypes and lower percentage of IgA and IgG isotypes in SSA+SjD participants (Figure 4A, Supplemental Table 6), consistent with more early forms of B cells and less progression to memory cells (Supplemental Figure 7A-C). Further, within transitional B cells, SSA+SjD samples had higher IgM and lower IgD percentages was lower compared to the two other groups (Figure 4B-C). This lower level of IgD suggests a potential bias toward marginal zone B cell differentiation, a delay in transitional B cell development, or premature inappropriate activation.

**Figure 4.**
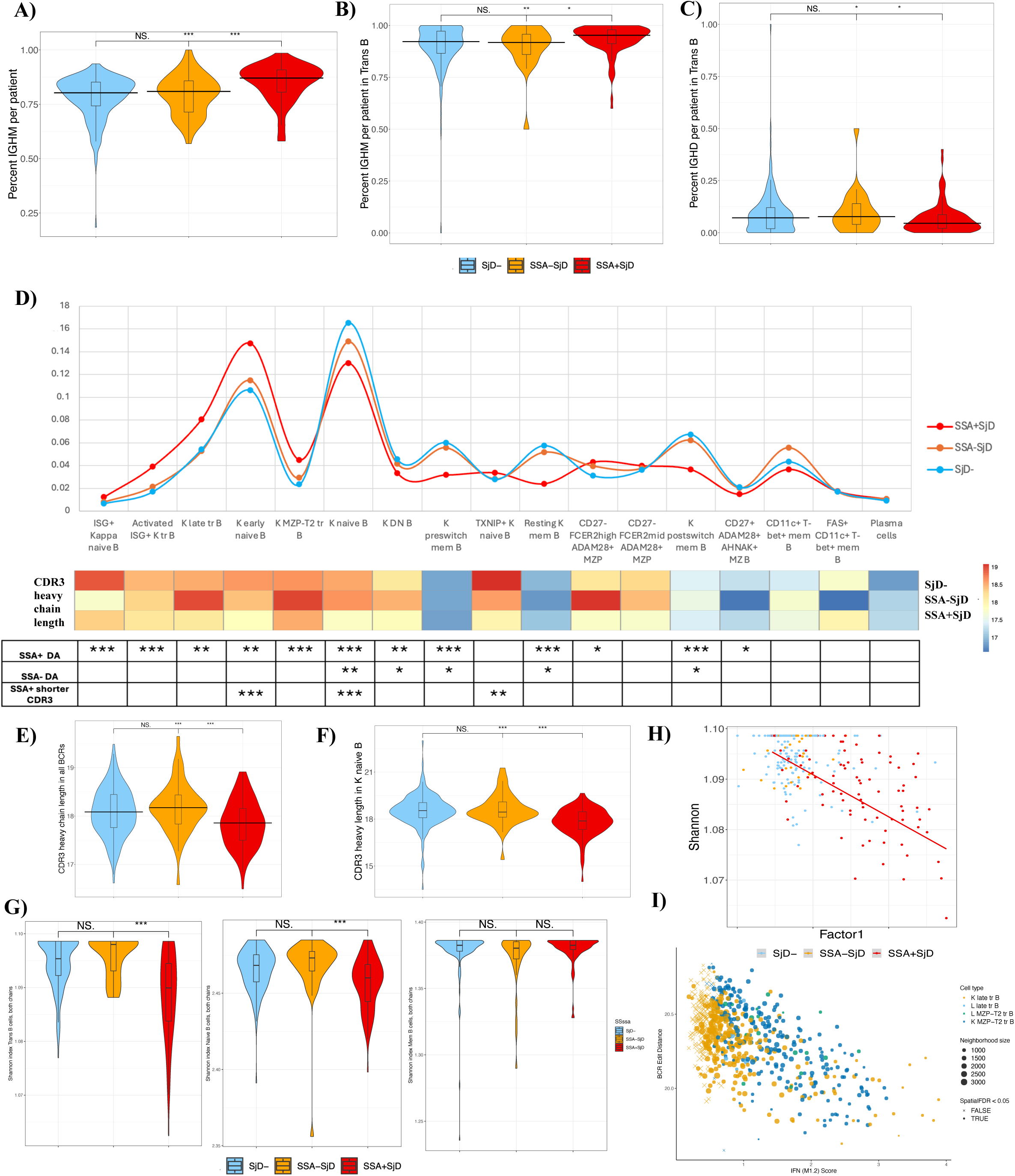
Cells of SSA+SjD patients have more transitional B cells, shorter heavy chains of BCRs, and less BCR diversity. A-C) mean BCR isotype proportions by SjD/SSA group: A) IgM in all cells, B) IgM in transitional B, C) IgD in transitional B; D) abundance of kappa cell types predicted by Dirichlet regression and ordered by pseudotime, with below CDR3 heavy chain length and significance of DA and CDR3 heavy chain length differences by cell type (***FDR p < 1e-5, **FDR p<0.001, *FDR p<0.05 from Dirichlet regression and t-test, respectively); E-F) mean BCR CDR3 heavy chain length per patient by SjD/SSA group in E) all cells and F) kappa naïve B cells; G) Shannon diversity index for transitional B cells; naïve B cells, and memory B cells; H) BCR Shannon index by Factor 1 in transitional B cells; I) Decreasing BCR similarity (edit distance) with increased IFN score (M1.2) in Milo transitional B neighborhoods.

Differential abundance analysis across all B cells illustrates reinforced transition from transitional B cells to marginal zone B precursors compared to follicular pathway in SSA+SjD participant samples (Figure 4D for predicted abundance for Kappa light chain-expressing B cells of the three groups along a pseudotime trajectory confirmed via Monocle^32^ inference; Supplemental Figure 7D for Lambda light chain-expressing B cells).

We also observed a significantly lower CDR3 heavy chain length overall in SSA+SjD patients (p=0.00015 versus SjD-, p=0.00035 versus SSA-SjD, Figure 4E). Within cell clusters, there was a trend to shorter CDR3 heavy chains in both Kappa and Lambda transitional and naive cells, with significantly shorter chains in Kappa naive B, Kappa early naive, and TXNIP+ K naive B cells (Figures 4D, 4F, Supplemental Table 7). B cell diversity evaluated by the Shannon index clearly demonstrated a loss of diversity in SSA+SjD samples compared to the other two groups, but this reduction was restricted to early B cell forms, especially transitional as well as naive B cells (Figure 4G). This loss of diversity among transitional B cells was inversely proportional to both the intensity of the IFN score (Figure 4H, r=-0.58, p=2e-9) and loss of BCR heavy chain length (r=0.25, p=0.017). Finally, we saw decreasing BCR similarity (edit distances) in transitional and naive B cell neighborhoods as IFN M1.2 increased (Figure 4I).

Two heavy-chain variable (IGHV) genes had significantly higher proportions in SSA+SjD versus SjD- groups (Supplemental Figure 7E-F), IGHV3-13 and IGHV4-34, specifically in transitional and naive subsets (Supplemental Table 8). IGHV4-34, known to be strongly associated with autoreactivity ^33,34^, was notably enriched in nearly all transitional B cells, including marginal zone precursors (MZP) of the extrafollicular pathway.

### Transitional B cells: key drivers of inflammation and autoimmunity

Transitional B cells in the SSA+SjD group are far from passive players. Instead, they are central to a network driving inflammation and sustaining autoreactive populations. These cells show enhanced communication, both as senders and receivers, with EM CD4+ and CD8+ T cells, naive CD8+ T cells, and classical monocytes, compared to other groups (Figure 5A-C). Further analysis of communication using refined clusters revealed that in SSA+SjD, transitional B cells primarily interact with terminally differentiated T cells (TEMRA CD8+ T, ICOShigh CCR4+ EM CD4+ T), NK cells, and IFN-I-stimulated populations (ISG+ monocytes) (Figure 5D). Key outgoing signals from transitional B cells include CD23, supporting survival and maturation; IL-16, a CD4+ T cell chemoattractant; ADGRE, promoting cell migration; and CypA, both an inflammation promoter (via NF-kB and MAPK) and a strong inhibitor of apoptosis (Figure 5E) ^35^. Key incoming signaling pathways include CD45, crucial for BCR signaling^36^; BAFF, B cell activation factor; and strong MIF signals (Figure 5F-G) promoting activation, HLA-II presentation through CD74, and migration signals via CXCR4 from bone marrow to blood ^33^ or from blood to lymphoid organs and inflamed tissues ^37^. HLA gene upregulation in SSA+SjD transitional B cells supported this (Figure 2I), with cell communication analysis revealing reinforced pathways involving HLA: a triangulation between classical monocytes and EM CD8+ T cells via HLA-I, and between classical monocytes and EM CD4+ T cells via HLA-II (Figure 5H-I). Communication with CD16bright NK cells through CLEC2-KLRB1 is reinforced, potentially enhancing NK cytotoxicity, but could also limit transitional B cell anergy (Figure 5J-K) ^38^. These dynamics contribute to sustained immune activation and support the persistence of autoreactive B cells.

**Figure 5.**
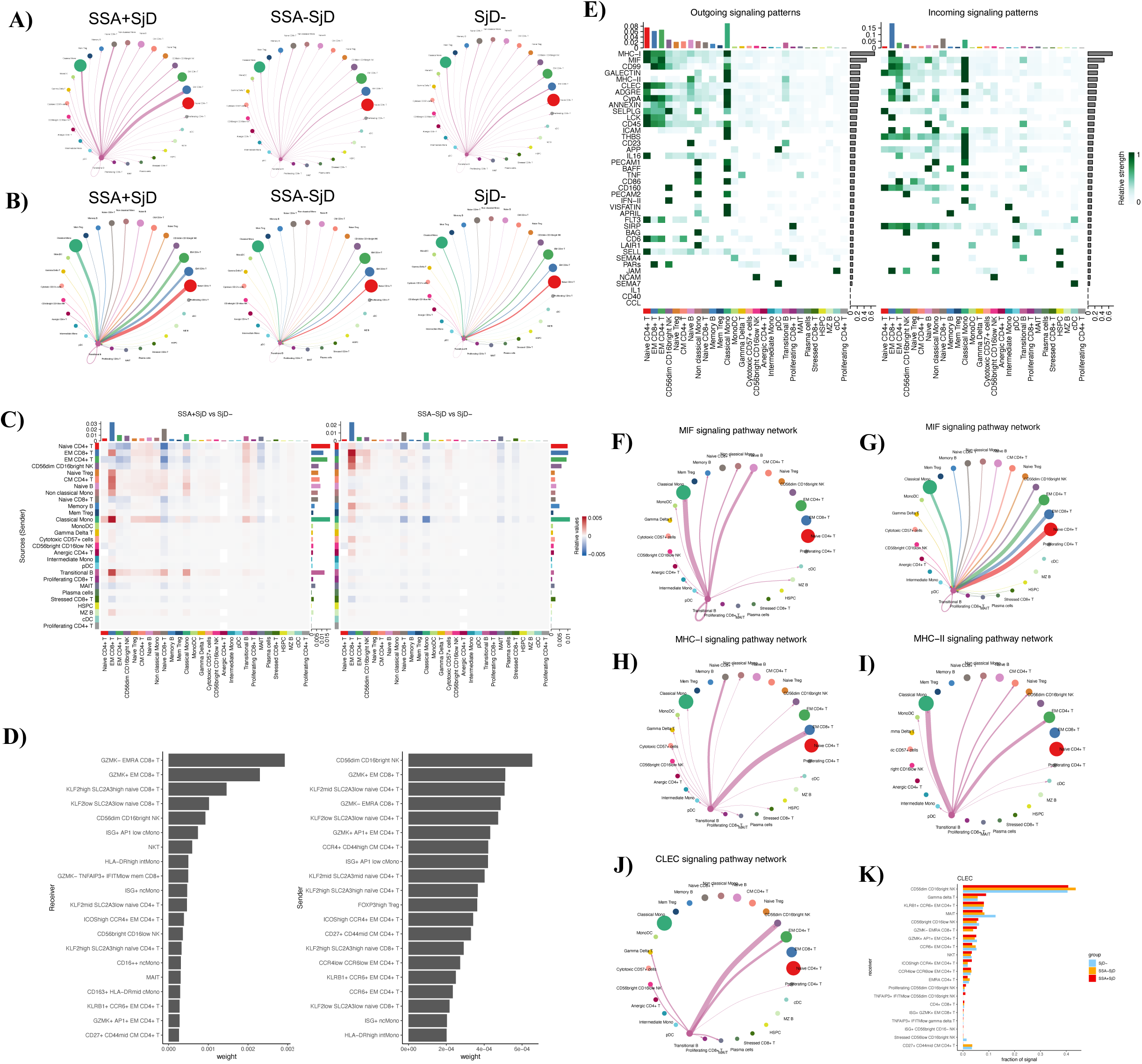
Transitional B cells communicate with other cell types, driving inflammation and autoimmunity. Outgoing (A) and incoming (B) signaling strength between Transitional B and other cell types. C) Heatmaps showing differential signaling strength between SSA+SjD and SjD- (left), and SSA-SjD and SjD- (right) across cell type sender/receiver pairs. D) Top outgoing (left) and incoming (right) fine annotations across all pathways for Transitional B in SSA+SjD. E) Heatmaps of outgoing (left) and incoming (right) signaling strength by pathway. Circle plots showing Transitional B signaling in SSA+SjD for MIF outgoing (F) and incoming (G); and MHC-I (H), MHC-II (I), and CLEC (J) outgoing. K) Top weighted receiver fine annotations for CLEC, plotted by fraction of signal in each group.

## Discussion

SjD is a complex and heterogeneous autoimmune disorder that significantly impacts patients’ quality of life ^39,40^. Advancing our understanding of its immunopathogenesis is critical for improving early diagnosis, refining severity stratification, and developing targeted therapies with the potential to modify disease progression ^19^. However, the substantial heterogeneity of SjD presents major challenges for both diagnosis and treatment, underscoring the need for a more precise disease characterization.

SjD stands out among autoimmune connective tissue diseases as a unique case where the hallmark serological biomarker, anti-SSA antibodies, is absent in nearly half of patients. In this study, we investigated this clinical dichotomy through a comprehensive single-cell transcriptomic analysis of SjD in a large cohort of 423 samples, uncovering profound molecular distinctions between these subgroups. Our findings reveal that anti-SSA antibodies are more than just a clinical marker, they represent a key molecular discriminator, defining distinct pathogenic pathways that drive disease heterogeneity. Interestingly, anti-SSA antibodies strongly correlated with the IFN score, whereas EGM presence did not, suggesting that these antibodies are deeply intertwined with the IFN pathway rather than merely reflecting the systemic clinical spread of SjD. To date, most SjD studies have combined SSA-positive and SSA-negative participants, leaving their distinct differences underexplored. A few studies have highlighted key genomic and immunological distinctions, including the association of anti-SSA antibody with HLA class II genes ^41–43^ (specifically HLA-DQB1-0201 and HLA-DRB1-0301^44^), as well as other immune-related polymorphisms (*HCP5* ^45^, *GTF2I* ^46^, and *CD5*^47^), and upregulation of ISGs but these findings have been limited by bulk transcriptomic approaches ^17,18^ and small sample sizes ^48^. Our scRNA-seq analysis of PBMCs provides an unprecedented level of resolution into the immune landscape of SjD. We uncovered a dominant and persistent IFN-I signature exclusively linked to SSA-positive individuals, further reinforcing the notion that SjD encompasses distinct immunological subtypes. Notably, the absence of ISG upregulation in SSA-negative patients challenges the prevailing pathogenic hypothesis, which posits that IFN-I plays a central role in recruiting immune cells to the exocrine glands and driving lymphocytic infiltration ^8,9^. This is further supported by recent transcriptomic ^49^ and metabolomic ^50^ studies of salivary glands, which reveal striking molecular differences between SSA-positive and SSA-negative patients. Collectively, these findings establish that SjD should be viewed as two distinct disease entities: one systemic, driven by anti-SSA antibodies and IFN-I activation, and another localized, primarily affecting the exocrine glands without systemic immune dysregulation.

In addition to defining these immunological subtypes, our study provides strong evidence that IFN-I stimulation profoundly alters immune cell maturation, particularly in B cells. Across all major immune cell types—T/NK cells, monocytes, and B cells—we observed disrupted subpopulation distributions, with B cell maturation being the most affected. Specifically, IFN-I signaling accelerates the early maturation of transitional B cells, the circulating precursors of mature naïve B cells. Transitional B cells play a critical role in immune tolerance, serving as a checkpoint where autoreactive clones are typically eliminated through clonal deletion, anergy, or receptor editing. Previous studies have reported an increased proportion of transitional B cells in SjD ^51–53^, but these analyses were limited in power and primarily defined transitional B cells based on surface markers (CD27-CD24highCD38high), mainly capturing the T2 stage of development. These findings, which suggested an impact on the extrafollicular pathway, lacked sufficient granularity to resolve the maturation trajectory of these cells. By integrating single-cell transcriptomics with surface protein profiling, our study demonstrates that IFN-I signaling alters all stages of transitional B cells, when defined by their pre-BCR and expression *VPREB3.* Additionally, it creates a pro-inflammatory environment that enables transitional B cells, through interactions with activated and proinflammatory cells, to bypass the normal process of anergy. This dysregulation promotes the survival of autoreactive B cells in both follicular and extrafollicular pathways, a process further supported by our observation of naive B cells with reduced BCR diversity and shorter CDR3 regions in SSA-positive patients. These findings suggest a fundamental shift in B cell development under IFN-I influence, creating an optimal environment for the persistence of autoreactive clones. However, whether this dysregulation originates peripherally, by promoting the survival of transitional B cells, or centrally, within the bone marrow at the precursor stage, remains an open question.

A major strength of our study is its unprecedented level of granularity, achieved by combining scRNA-seq with surface protein characterization in a dataset of over 1.5 million cells. This approach provides a comprehensive understanding of immune dysregulation in SjD, particularly in transitional B cells under IFN-I influence. Another key strength is the comparison of SjD participants to non-SjD symptomatic controls rather than healthy individuals. Non-SjD symptomatic controls are more closely matched in terms of age, sex distribution, medication use, and other confounding factors, ensuring that our findings reflect disease-specific immune alterations rather than general differences between patients and healthy subjects. The main limitation of our study is the lack of direct mechanistic experimentation, which prevents us from definitively establishing a causal relationship between IFN-I signaling, B cell dysregulation, and anti-SSA antibody production. Future studies should focus on functional validation of these pathways to determine whether IFN-I blockade can restore normal B cell maturation and prevent autoreactive clones from emerging.

In summary, our study represents the most comprehensive single-cell analysis of peripheral immune dysregulation in SjD to date. We identify a dominant IFN-I signature in SSA+SjD that correlates with profound B cell dysregulation and supports a model of two immunologically distinct disease subtypes. These findings challenge the traditional classification of SjD and provide a strong rationale for biomarker-driven therapeutic strategies targeting IFN-I signaling and B cell maturation. Future studies should explore targeted interventions in these molecularly defined subgroups to assess their potential for altering disease progression and improving patient outcomes.

## Materials and methods

### Ethics

This study was approved by the Institutional Review Board of the University of California, San Francisco (UCSF) (#10-02551), and was conducted in compliance with the Declaration of Helsinki. All participants signed a written informed consent.

### Study population and specimen collection

Among the 3,514 participants from the previously-described SICCA cohort ^54^, we selected participants among those with available stored PBMCs (with at least 2 vials in the SICCA biorepository), all of whom were recruited at the UCSF site center. To ensure representation from specific subgroups, we randomly selected participants from within these sub-groups. The eligibility criteria for the SICCA cohort, which included signs and symptoms suggestive of SjD among women and men aged 21 years and older, have been previously described ^3,54^. We excluded participants with 1) other connective tissue diseases (systemic lupus erythematosus, rheumatoid arthritis, immune myopathy, systemic scleroderma, and mixed connective tissue disease) based on physician confirmed diagnosis; 2) indeterminate status, due to missing data among the five criteria items of the 2016 ACR-EULAR classification criteria for SjD. At the sequencing step, we also excluded samples for whom were unable to map cells to individuals based on genetics. A flow-chart of this sample selection process is in Supplemental Figure 1.

### Clinical and demographic data

We extracted data from the SICCA database pertaining to general demographic information, including age, sex, and self-reported ancestry, categorized as follows: African American, Asian, White, Latin American, Native American, and Pacific Islander.

Participants in the SICCA project had previously been evaluated for eye dryness with a Schirmer type I test without anesthesia, and with an ocular surface staining score (OSS), and for objective mouth dryness using a 5-minute unstimulated whole salivary (UWS) flow rate. Participants also had labial salivary gland (LSG) biopsy collected, with evaluation of focal lymphocytic sialadenitis (FLS) quantified by the focus score (FS) and for germinal centers, these were assessed by the same two pathologists, who were calibrated with respect to histopathologic interpretation of FLS and computation of the FS. The evaluation also included serologic immune assays with anti-SSA/SSB antibodies, antinuclear antibodies (ANA) and their titer, and measurements of rheumatoid factors (RF), immunoglobulins G (IgG), complement C3 and C4 fractions, all performed by the same commercial laboratory (Quest Diagnostics, Madison, NJ).

SjD was assessed according to the 2016 ACR/EULAR criteria ^3^. Participants not meeting these criteria were classified as non-SjD symptomatic participants and labeled as SjD-. Participants meeting the SjD classification criteria were labeled as SjD+ and further divided based on the presence or absence of anti-SSA antibodies as follows: SSA-SjD for those without detectable anti-SSA antibodies, and SSA+SjD for those testing positive for these antibodies. For specific analyses, the SSA+SjD group was divided based on the presence (SSB+SSA+SjD) or absence (SSB-SSA+SjD) of anti-SSB antibodies.

To have an overview of B-cell activity, we derived a binary variable, named B lymphocytes signs of activity (BLA), based on the presence of at least two of the following five markers: presence of rheumatoid factors, IgG level >2000 mg/dL, IgG level <500 mg/dL, decreased level of C3 (<90 mg/dL), and decreased level of C4 (<16 mg/dL) ^55^.

Because the SICCA project was initiated in 2003, which preceded the development and validation of the EULAR Sjögren Syndrome Disease Activity Index (ESSDAI)^55^, the ESSDAI could not be included among the variables measured as part of SICCA. However, we used the following clinical and biological SICCA variables inspired by the ESSDAI items to assess the presence/absence of extraglandular manifestations (EGM): history of autoimmune hemolytic anemia and/or anemia defined by hemoglobin level <120 g/L, history of immune thrombocytopenic purpura and/or thrombocytopenia defined by platelet level <150*10^9^/L, history of immune neutropenia and/or neutropenia defined by neutrophils level <1.5*10^9^/L, history of immune lymphopenia and/or lymphopenia defined by neutrophils level <1*10^9^/L, inflammatory arthralgia and/or arthritis, acute or chronic erythema multiforma-like lesions, cutaneous vasculitis, pleuritis, pericarditis, kidney involvement (glomerulonephritis, interstitial nephritis, renal tubular acidosis), central nervous system involvement, interstitial lung disease, splenomegaly, lymphadenopathy, and lymphoma.

Finally, we also considered data on the immune-modulating drugs that participants reported taking at the time of PBMC sampling, collected as follows: corticosteroids, antimetabolites, TNFα inhibitors, alkylating agents, anti-CD20 drugs taken within the past 12 months, other DMARDs, and hydroxychloroquine.

### PBMCs preparation

SICCA PBMCs were collected on acid–citrate–dextrose, isolated using Ficoll separation, and cryopreserved in the SICCA biorepository at UCSF. Frozen PBMCs from 433 samples including 93 for patients with follow-up (i.e. 340 for baseline) (Supplemental Figure 1) were processed into 28 pools across five processing batches; each pool was replicated in 4-5 libraries. PBMCs were thawed, transferred into RPMI1640 medium with L-glutamine and HEPES (Corning, Glendale, Arizona, USA) containing 10% of Fetal Calf Serum and warmed at 37°C, counted and viability-checked using a Cellaca MX High-throughput Automated Cell Counter (Nexcelom, Lawrence, Massachusetts, United States), resuspended, and pooled with 85,000 cells/pool. For batches 3, 4 and 5, pooled cells were resuspended in 45 µL of Cell Staining Buffer (BioLegend 420201, San Diego, California, USA) and incubated at 4°C for 10 minutes with 10% Human TruStain FcX (5 µL) (BioLegend 422301, San Diego, California, USA) to block non-specific staining. Then, we added in the latter solution 25 µL of a mix including TotalSeq-C Human antibodies Cocktail V1.0 (BioLegend 399905, San Diego, California, USA), TotalSeq-C anti-human CD197 (CCR7) (BioLegend 353251, San Diego, California, USA), and TotalSeq-C anti-human CD138 (Syndecan-1) (BioLegend 352327, San Diego, California, USA) completed with 25 µL Cell Staining Buffer (BioLegend 420201, San Diego, California, USA) for a 30-minute incubation at 4°C. Cells were then resuspended in 3 mL of PBS with 0.04% BSA and strained using a 40-micron cell strainer (Corning 352340, Glendale, Arizona, USA). ADT-labeled (batches 3 to 5) and unstained (batches 1 and 2) cells were loaded onto a 10X Chromium Controller.

### RNA, antibody-derived tags (ADT), and BCR/TCR library preparation, and sequencing

Samples underwent single-cell encapsulation and cDNA library preparation using the Chromium Single Cell 5′ v1.1 Reagent Kits (10× Genomics, Pleasanton, California, USA). scRNA-seq libraries were prepared following the protocol provided by the 10x Genomics Chromium NEXTGEM single-cell V(D)J version 1.1. Cell suspensions were loaded into a 10x Genomics Chromium Controller instrument to generate single-cell gel bead-in emulsions (GEMs). The term “GEM” refers to the mixture of cells, reagents, and gel beads containing barcodes wrapped in oil droplets. As the gel beads in the GEM dissolved, cells were lysed to release mRNA through reverse transcription. After reverse transcription step, DynaBeads were used to purify the barcoded cDNA, and cycles of PCR amplification were then performed. The amount of resulting amplified cDNA was used to construct TCR/BCR-enriched libraries and 5′ Gene Expression libraries. The cDNA was fragmented, performed double size selection with SPRI beads, followed with adaptors ligation and sample index PCR. The 5’ Gene Expression libraries contain the P5 and P7 are constructed. For BCR/TCR, full-length V(D)J segments were enriched from amplified cDNA, the enriched TCR and Ig transcripts went through subsequent fragmentation, ligation, and sample index PCR, to construct the TCR and BCR library. The libraries were sequenced on an Illumina Novaseq 6000. The sequencing cycles were at 28 cycles for R1 and 90 for R2.

### Single-cell RNA-seq data preprocessing

Sequencing data were processed using the 10x Genomics Cell Ranger version 6.0.2. Fastq files were generated from Ilumina bcl files with the Cell Ranger subroutine *mkfastq*. Fastq files were then processed with Cell Ranger’s *count* to align RNA reads against 10x Reference 2020-A (GENCODE v32/Ensembl 98). Redundant unique molecular identifiers (UMI) were filtered, and a gene–cell barcode matrix was generated with *count*. *Mkfastq* and *count* were run with default parameters.

Freemuxlet ^15^ was used to identify genotype clusters for each library, resulting in a total of 2,366,418 singlets. Freemuxlet clusters were compared to previously collected and processed and genotyping data, and significant (p < 0.05) cluster-genotype pairings were retained, resulting in identification of 423 out of 434 individual samples (including baseline and follow-up samples). Freemuxlet clusters from unassigned individuals were removed, along with 2 individuals not in the cohort. Additional removal of doublets with the same genotype was performed using DoubletFinder ^56^ (58,362 droplets, overall intra-doublet rate 2.5%), with per-library intra-individual doublet rates estimated based on the number of inter-individual doublets from Freemuxlet and individuals in that library.

Genes were initially retained if they were expressed in at least 12 cells in each library. Cells were filtered to those having 500-1000 UMIs, 300-3000 total genes, 0-15% mitochondrial genes, 0-60% ribosomal genes, 3% or less platelet genes (*PPBP* or *PF4*), and 15% or less hemoglobin genes (prefix *HB* but not *HBP*). When merging libraries, we also required genes to be in at least 80% of libraries (92/115).

In three out of five batches, we additionally measured paired cell surface protein measurements via CITE-Seq (Antibody-derived tags (ADT)). The corresponding libraries for cell surface protein data from antibody-derived tags (ADT) were aligned with Cell Ranger, then normalized and background corrected with DSB ^57^. Normalized ADT data were integrated using Reciprocal Principal Component Analysis (RPCA) as the reduction method in the Seurat function *FindIntegrationAnchors()* ^58^.

BCR/TCR were aligned and initial filtering was performed with the CellRanger *vdj* function. Filtered contigs from CellRanger were input into scRepertoire ^59^ (version 1.10.0) to identify clonotypes. Additionally, in the case there were more than one heavy and light chain for BCR in a cell, scRepertoire retained chains with the largest UMIs were retained. Following clonotype identification, incomplete BCRs/TCRs (BCRs missing a heavy or light chain; TCRs missing an alpha or beta chain) were removed.

Finally, all RNA count matrices were log-normalized and merged into a single Seurat object for downstream analysis. After all preprocessing, QC, and merge of libraries we retained 1,537,678 cells with 16,524 genes for analysis. A standard Seurat pipeline was executed on the merged samples for log-normalization, identification of variable features (2000 via the “vst” method), and scaling ^58^. During scaling, percent mitochondrial genes, percent ribosomal genes, number of UMIs, and number of features were regressed out. Harmony ^60^ was used to adjust for pool membership. Batch-corrected protein (ADT) count data were then added as an additional assay to the processed Seurat object.

### Cell type annotations

Cell clustering and cell type annotations of clusters were done at the level of all cells, and then refined, by reclustering within the major cell type groups of B cells, T/NK cells, and Monocytes & Dendritic (MD) cells. Cell clustering was performed using the Louvain algorithm, a modularity optimization technique, with a stepwise increase in resolution granularity. Starting at a resolution of 0.25, increments of 0.25 were applied until the first non-individual-specific cluster comprising less than 0.01% of total cells was identified.

Gene and protein expression data from CITE-seq were both utilized to identify cell types. Differential expression analysis was conducted using Seurat *FindAllMarkers()* function filtering to genes or proteins with a log fold change of at least 0.25 and detected in at least 10% of cells in either population by comparing each cell type against all other clusters. The resulting identified differential genes and proteins were carefully analyzed for cell type identification and labeling.

Annotations were primarily performed using classical markers for cell subtypes, derived from both gene and surface protein expression data. For consistency, surface protein expressions were labeled using their corresponding gene names, regardless of whether they represent genes or proteins. Those markers are listed in the Supplemental Table 8. If multiple clusters retained the same label after initial tagging with classical markers, additional annotations were derived based on differentially expressed markers, evaluated for either their expression intensity or relevance. Marker relevance was assessed using the Human Protein Atlas database ^61^ and/or a thorough review of relevant literature. Special attention was given to the identification of ISG expressions. HSP genes were analyzed to identify stressed cells. Cells with high expression levels of *ACTG1, HIST1H4C, STMN1, HMGB2, TUBB, MKI67*, and *MT2A* were classified as proliferating cells. Conversely, cells with low expression levels across all genes and surface proteins were categorized as anergic cells.

### Differential abundance and gene expression analyses

We utilized Dirichlet regression ^28^ for DA analysis, as it handles compositionality and does not require a reference cell type. We adjusted for age, sex, and top four genetic PCs in this analysis. Abundance principal components were compared to phenotypes using Pearson correlation and across groups with t-tests.

We used EdgeR ^27^ for DE testing, adjusting for age, sex, number of expressed genes, top 4 genetic principal components for ancestry, and pool. We tested genes expressed in at least 5% of the cells in both of the two conditions under test (e.g. SSA+SjD vs SjD-). Gene set enrichment analysis was performed with the enrichGO function from ClusterProfiler ^62^ based on the ordered logFC from DEG analysis, with the GO biological process ontology as the gene set reference. P-values were Benjamini-Hochberg corrected, and cutoff at adjusted-p < 0.05.

### Analysis of multicellular transcriptional program

We used Single-cell Interpretable Tensor Decomposition ^29^ (scITD) to examine expression patterns across multiple cell types that are correlated with SjD and its subphenotypes. This method produces factors which have gene by cell type loadings, reflecting the contribution of genes in each cell type to the pattern, and participant weights (“donor scores”), reflecting the degree of each factor found within the individual participant. As scITD excludes patients having zero cells of any included clusters, we removed clusters present in the fewest patients, namely Anergic CD4+T cells, Proliferating CD4+T cells, HSPCs, Plasma cells, and cDCs; 302 participants had at least one of the remaining cell types and were included in the main analysis. We chose “rank” parameters of 12 factors and 24 gene sets, basing this selection on scITD-provided functions to evaluate reconstruction error and factor stability at different rank levels.

### Interferon signature

To compare the Factor 1 donor score obtained from scITD to IFN signatures at the patient level, we first computed pseudobulk gene expression across all cells for each patient. We then summed the expression of genes in each of the modules namely M1.2 (*IFI44, IFI44L, IFIT1, IFIT3*, and *MX1*), M3.4 (*ZBP1, EIF2AK2, IFIH1, PARP9,* and *GBP4*), and M5.12 (*PSMB9, NCOA7, TAP1, ISG20*, and *SP140*) ^30^. We compared these scores with each other and with SjD phenotypes (such as anti-SSA and anti-SSB) using Spearman correlation.

### Cell trajectories and neighborhoods

Trajectory analysis was performed with Monocle ^32^ on the kappa and lambda B cell subpopulations separately. Clustering and pseudotime analysis were performed on the Harmony-adjusted UMAP (described above), with root cells set to a randomly sampled subset of 1000 cells in the transitional B population.

MiloR ^31^ was used for nearest neighbor based differential expression analysis with k=500 and d=30. The resulting neighborhoods were then annotated by their majority cell subtype, neighborhoods with less than 30% of any cell subtype were annotated as “Mixed”. The resulting milo neighborhoods were tested for association with SSA/SjD status using a similar framework to DE analysis (adjusting for age, sex, batch, and four ancestry PCs). Contrasts for SSA+SjD vs SjD- and SSA-SjD vs SjD- were extracted from the model, and the results filtered to neighborhoods with spatial FDR < 0.05.

### BCR/TCR analysis

After scRepertoire pre-processing, unique clonotypes per patient were retained for downstream analysis. Isotype proportions were determined as the percentage of unique clonotypes containing each isotype (e.g. IgM) per patient. Statistical comparisons between SSA+SjD, SSA-SjD, and SjD- were done using Wilcoxon tests (isotype and V-gene proportions) and t-tests (CDR3 length).

BCR diversity using all clonotypes was measured per patient using the Shannon Index produced by the *clonalDiversity()* function of scRepertoire. Two outliers were removed, both of whom had immune cytopenias. We also filtered to remove individuals with the 15% fewest BCRs for this analysis, as scRepertoire performs bootstrap sampling of the smallest sample size to avoid bias and this can result variances too low for statistics. Levenshtein edit distance (from rapidfuzz package^63^) was also used to examine differences between BCRs.

### Cell-cell Communication Analysis

Cell-cell communication analysis was performed using CellChat (v2.1.2)^64^ separately for each of the three groups (SSA+SjD, SSA-SjD, SjD-), and the results compared. Analysis was performed twice, with first the original and then reclustered annotations.

### Software and packages

All analyses were performed in R version v4.2.0 (R Core Team 2021) with Seurat v4.1.1 ^65^.

## References

1. Mariette, X., and Criswell, L.A. (2018). Primary Sjögren’s Syndrome. N. Engl. J. Med. 378, 931–939. 10.1056/NEJMcp1702514.

2. Ansari, N., and Salesi, M. (2024). The association between primary Sjogren’s syndrome and non-Hodgkin’s lymphoma: a systematic review and meta-analysis of cohort studies. Clin. Rheumatol. 43, 2177–2186. 10.1007/s10067-024-06993-6.

3. Shiboski, C.H., Shiboski, S.C., Seror, R., Criswell, L.A., Labetoulle, M., Lietman, T.M., Rasmussen, A., Scofield, H., Vitali, C., Bowman, S.J., et al. (2017). 2016 American College of Rheumatology/European League Against Rheumatism classification criteria for primary Sjögren’s syndrome: A consensus and data-driven methodology involving three international patient cohorts. Ann. Rheum. Dis. 76, 9–16. 10.1136/annrheumdis-2016-210571.

4. Daniels, T.E., Criswell, L.A., Shiboski, C., Shiboski, S., Lanfranchi, H., Dong, Y., Schiødt, M., Umehara, H., Sugai, S., Challacombe, S., et al. (2009). An early view of the international Sjögren’s syndrome registry. Arthritis Rheum. 61, 711–714. 10.1002/art.24397.

5. Brito-Zerón, P., Acar-Denizli, N., Ng, W.-F., Zeher, M., Rasmussen, A., Mandl, T., Seror, R., Li, X., Baldini, C., Gottenberg, J.-E., et al. (2018). How immunological profile drives clinical phenotype of primary Sjögren’s syndrome at diagnosis: analysis of 10,500 patients (Sjögren Big Data Project). Clin. Exp. Rheumatol. 36 *Suppl 112*, 102–112.

6. Urbanski, G., Gury, A., Jeannin, P., Chevailler, A., Lozac’h, P., Reynier, P., Lavigne, C., Lacout, C., and Vinatier, E. (2022). Discordant Predictions of Extraglandular Involvement in Primary Sjögren’s Syndrome According to the Anti-SSA/Ro60 Antibodies Detection Assay in a Cohort Study. J. Clin. Med. 11, 242. 10.3390/jcm11010242.

7. Martel, C., Gondran, G., Launay, D., Lalloué, F., Palat, S., Lambert, M., Ly, K., Loustaud-Ratti, V., Bezanahary, H., Hachulla, E., et al. (2011). Active immunological profile is associated with systemic Sjögren’s syndrome. J. Clin. Immunol. 31, 840–847. 10.1007/s10875-011-9553-3.

8. Verstappen, G.M., Pringle, S., Bootsma, H., and Kroese, F.G.M. (2021). Epithelial-immune cell interplay in primary Sjögren syndrome salivary gland pathogenesis. Nat. Rev. Rheumatol. 17, 333–348. 10.1038/s41584-021-00605-2.

9. 9. Baldini, C., Fulvio, G., La Rocca, G., and Ferro, F. (2024). Update on the pathophysiology and treatment of primary Sjögren syndrome. Nat. Rev. Rheumatol. 20, 473–491. 10.1038/s41584-024-01135-3.

10. Zheng, L., Zhang, Z., Yu, C., and Yang, C. (2010). Expression of Toll-like receptors 7, 8, and 9 in primary Sjögren’s syndrome. Oral Surg. Oral Med. Oral Pathol. Oral Radiol. Endod. 109, 844-850. 10.1016/j.tripleo.2010.01.006.

11. Aqrawi, L.A., Jensen, J.L., Øijordsbakken, G., Ruus, A.-K., Nygård, S., Holden, M., Jonsson, R., Galtung, H.K., and Skarstein, K. (2018). Signalling pathways identified in salivary glands from primary Sjögren’s syndrome patients reveal enhanced adipose tissue development. Autoimmunity 51, 135–146. 10.1080/08916934.2018.1446525.

12. Lee, J., Lee, J., Kwok, S.-K., Baek, S., Jang, S.G., Hong, S.-M., Min, J.-W., Choi, S.S., Lee, J., Cho, M.-L., et al. (2018). JAK-1 Inhibition Suppresses Interferon-Induced BAFF Production in Human Salivary Gland: Potential Therapeutic Strategy for Primary Sjögren’s Syndrome. Arthritis Rheumatol. Hoboken NJ 70, 2057–2066. 10.1002/art.40589.

13. Zhao, T., Zhang, R., Li, Z., Qin, D., and Wang, X. (2024). A comprehensive review of Sjögren’s syndrome: Classification criteria, risk factors, and signaling pathways. Heliyon 10, e36220. 10.1016/j.heliyon.2024.e36220.

14. Lessard, C.J., Li, H., Adrianto, I., Ice, J.A., Rasmussen, A., Grundahl, K.M., Kelly, J.A., Dozmorov, M.G., Miceli-Richard, C., Bowman, S., et al. (2013). Variants at multiple loci implicated in both innate and adaptive immune responses are associated with Sjögren’s syndrome. Nat. Genet. 45, 1284–1292. 10.1038/ng.2792.

15. Taylor, K.E., Wong, Q., Levine, D.M., McHugh, C., Laurie, C., Doheny, K., Lam, M.Y., Baer, A.N., Challacombe, S., Lanfranchi, H., et al. (2017). Genome-Wide Association Analysis Reveals Genetic Heterogeneity of Sjögren’s Syndrome According to Ancestry. Arthritis Rheumatol. Hoboken NJ 69, 1294–1305. 10.1002/art.40040.

16. 16. Bodewes, I.L.A., Al-Ali, S., van Helden-Meeuwsen, C.G., Maria, N.I., Tarn, J., Lendrem, D.W., Schreurs, M.W.J., Steenwijk, E.C., van Daele, P.L.A., Both, T., et al. (2018). Systemic interferon type I and type II signatures in primary Sjögren’s syndrome reveal differences in biological disease activity. Rheumatol. Oxf. Engl. 57, 921–930. 10.1093/rheumatology/kex490.

17. Trutschel, D., Bost, P., Mariette, X., Bondet, V., Llibre, A., Posseme, C., Charbit, B., Thorball, C.W., Jonsson, R., Lessard, C.J., et al. (2022). Variability of Primary Sjögren’s Syndrome Is Driven by Interferon-α and Interferon-α Blood Levels Are Associated With the Class II HLA-DQ Locus. Arthritis Rheumatol. Hoboken NJ 74, 1991–2002. 10.1002/art.42265.

18. Hall, J.C., Baer, A.N., Shah, A.A., Criswell, L.A., Shiboski, C.H., Rosen, A., and Casciola-Rosen, L. (2015). Molecular Subsetting of Interferon Pathways in Sjogren’s syndrom. Arthritis Rheumatol. Hoboken NJ 67, 2437. 10.1002/art.39204.

19. Wang, B., Chen, S., Zheng, Q., Li, Y., Zhang, X., Xuan, J., Liu, Y., and Shi, G. (2021). Early diagnosis and treatment for Sjögren’s syndrome: current challenges, redefined disease stages and future prospects. J. Autoimmun. 117, 102590. 10.1016/j.jaut.2020.102590.

20. Hou, X., Hong, X., Ou, M., Meng, S., Wang, T., Liao, S., He, J., Yu, H., Liu, L., Yin, L., et al. (2022). Analysis of Gene Expression and TCR/B Cell Receptor Profiling of Immune Cells in Primary Sjögren’s Syndrome by Single-Cell Sequencing. J. Immunol. Baltim. Md 1950 *209*, 238–249. 10.4049/jimmunol.2100803.

21. He, Y., Chen, R., Zhang, M., Wang, B., Liao, Z., Shi, G., and Li, Y. (2022). Abnormal Changes of Monocyte Subsets in Patients With Sjögren’s Syndrome. Front. Immunol. 13, 864920. 10.3389/fimmu.2022.864920.

22. Liu, J., Gao, H., Li, C., Zhu, F., Wang, M., Xu, Y., and Wu, B. (2022). Expression and regulatory characteristics of peripheral blood immune cells in primary Sjögren’s syndrome patients using single-cell transcriptomic. iScience 25, 105509. 10.1016/j.isci.2022.105509.

23. Zong, Y., Yang, Y., Zhao, J., Li, L., Luo, D., Hu, J., Gao, Y., Wei, L., Li, N., and Jiang, L. (2023). Characterisation of macrophage infiltration and polarisation based on integrated transcriptomic and histological analyses in Primary Sjögren’s syndrome. Front. Immunol. 14. 10.3389/fimmu.2023.1292146.

24. Xu, T., Zhu, H.-X., You, X., Ma, J.-F., Li, X., Luo, P.-Y., Li, Y., Lian, Z.-X., and Gao, C.-Y. (2023). Single-cell profiling reveals pathogenic role and differentiation trajectory of granzyme K^+^CD8^+^ T cells in primary Sjögren’s syndrome. JCI Insight 8. 10.1172/jci.insight.167490.

25. Gupta, S., Yamada, E., Nakamura, H., Perez, P., Pranzatelli, T.J., Dominick, K., Jang, S.-I., Abed, M., Martin, D., Burbelo, P., et al. (2024). Inhibition of JAK-STAT pathway corrects salivary gland inflammation and interferon driven immune activation in Sjögren’s disease. Ann. Rheum. Dis. 83, 1034–1047. 10.1136/ard-2023-224842.

26. Arvidsson, G., Czarnewski, P., Johansson, A., Raine, A., Imgenberg-Kreuz, J., Nordlund, J., Nordmark, G., and Syvänen, A.-C. (2024). Multimodal Single-Cell Sequencing of B Cells in Primary Sjögren’s Syndrome. Arthritis Rheumatol. 76, 255–267. 10.1002/art.42683.

27. Robinson, M.D., McCarthy, D.J., and Smyth, G.K. (2010). edgeR: a Bioconductor package for differential expression analysis of digital gene expression data. Bioinformatics 26, 139–140. 10.1093/bioinformatics/btp616.

28. Malawsky, D., and Gershon, T.R. (2023). scRNA-seq for Microcephaly Research [IV]: Dirichlet Regression for Single-Cell Population Differences. Methods Mol. Biol. Clifton NJ 2583, 123–125. 10.1007/978-1-0716-2752-5_11.

29. Mitchel, J., Gordon, M.G., Perez, R.K., Biederstedt, E., Bueno, R., Ye, C.J., and Kharchenko, P.V. (2024). Coordinated, multicellular patterns of transcriptional variation that stratify patient cohorts are revealed by tensor decomposition. Nat. Biotechnol. 10.1038/s41587-024-02411-z.

30. Chiche, L., Jourde-Chiche, N., Whalen, E., Presnell, S., Gersuk, V., Dang, K., Anguiano, E., Quinn, C., Burtey, S., Berland, Y., et al. (2014). Modular transcriptional repertoire analyses of adults with systemic lupus erythematosus reveal distinct type I and type II interferon signatures. Arthritis Rheumatol. Hoboken NJ 66, 1583–1595. 10.1002/art.38628.

31. Dann, E., Henderson, N.C., Teichmann, S.A., Morgan, M.D., and Marioni, J.C. (2022). Differential abundance testing on single-cell data using k-nearest neighbor graphs. Nat. Biotechnol. 40, 245–253. 10.1038/s41587-021-01033-z.

32. Trapnell, C., Cacchiarelli, D., Grimsby, J., Pokharel, P., Li, S., Morse, M., Lennon, N.J., Livak, K.J., Mikkelsen, T.S., and Rinn, J.L. (2014). The dynamics and regulators of cell fate decisions are revealed by pseudotemporal ordering of single cells. Nat. Biotechnol. 32, 381–386. 10.1038/nbt.2859.

33. Pugh-Bernard, A.E., Silverman, G.J., Cappione, A.J., Villano, M.E., Ryan, D.H., Insel, R.A., and Sanz, I. (2001). Regulation of inherently autoreactive VH4-34 B cells in the maintenance of human B cell tolerance. J. Clin. Invest. 108, 1061–1070. 10.1172/JCI12462.

34. Cappione, A., Anolik, J.H., Pugh-Bernard, A., Barnard, J., Dutcher, P., Silverman, G., and Sanz, I. (2005). Germinal center exclusion of autoreactive B cells is defective in human systemic lupus erythematosus. J. Clin. Invest. 115, 3205–3216. 10.1172/JCI24179.

35. Chen, L., Zeng, Z., Luo, H., Xiao, H., and Zeng, Y. (2024). The effects of CypA on apoptosis: potential target for the treatment of diseases. Appl. Microbiol. Biotechnol. 108, 28. 10.1007/s00253-023-12860-2.

36. 36. Pelanda, R., Greaves, S.A., Alves da Costa, T., Cedrone, L.M., Campbell, M.L., and Torres, R.M. (2022). B-cell intrinsic and extrinsic signals that regulate central tolerance of mouse and human B cells. Immunol. Rev. 307, 12–26. 10.1111/imr.13062.

37. Klasen, C., Ohl, K., Sternkopf, M., Shachar, I., Schmitz, C., Heussen, N., Hobeika, E., Levit-Zerdoun, E., Tenbrock, K., Reth, M., et al. (2014). MIF Promotes B Cell Chemotaxis through the Receptors CXCR4 and CD74 and ZAP-70 Signaling. J. Immunol. 192, 5273–5284. 10.4049/jimmunol.1302209.

38. Kirkham, C.L., and Carlyle, J.R. (2014). Complexity and Diversity of the NKR-P1:Clr (Klrb1:Clec2) Recognition Systems. Front. Immunol. 5, 214. 10.3389/fimmu.2014.00214.

39. Singh, A.G., Singh, S., and Matteson, E.L. (2016). Rate, risk factors and causes of mortality in patients with Sjögren’s syndrome: a systematic review and meta-analysis of cohort studies. Rheumatol. Oxf. Engl. 55, 450–460. 10.1093/rheumatology/kev354.

40. Huang, H., Xie, W., Geng, Y., Fan, Y., and Zhang, Z. (2021). Mortality in patients with primary Sjögren’s syndrome: a systematic review and meta-analysis. Rheumatol. Oxf. Engl. 60, 4029–4038. 10.1093/rheumatology/keab364.

41. Gottenberg, J.-E., Busson, M., Loiseau, P., Cohen-Solal, J., Lepage, V., Charron, D., Sibilia, J., and Mariette, X. (2003). In primary Sjögren’s syndrome, HLA class II is associated exclusively with autoantibody production and spreading of the autoimmune response. Arthritis Rheum. 48, 2240–2245. 10.1002/art.11103.

42. Barturen, G., Babaei, S., Català-Moll, F., Martínez-Bueno, M., Makowska, Z., Martorell-Marugán, J., Carmona-Sáez, P., Toro-Domínguez, D., Carnero-Montoro, E., Teruel, M., et al. (2021). Integrative Analysis Reveals a Molecular Stratification of Systemic Autoimmune Diseases. Arthritis Rheumatol. Hoboken NJ 73, 1073–1085. 10.1002/art.41610.

43. Thorlacius, G.E., Hultin-Rosenberg, L., Sandling, J.K., Bianchi, M., Imgenberg-Kreuz, J., Pucholt, P., Theander, E., Kvarnström, M., Forsblad-d’Elia, H., Bucher, S.M., et al. (2021). Genetic and clinical basis for two distinct subtypes of primary Sjögren’s syndrome. Rheumatol. Oxf. Engl. 60, 837–848. 10.1093/rheumatology/keaa367.

44. Teruel, M., Barturen, G., Martínez-Bueno, M., Castellini-Pérez, O., Barroso-Gil, M., Povedano, E., Kerick, M., Català-Moll, F., Makowska, Z., Buttgereit, A., et al. (2021). Integrative epigenomics in Sjögreńs syndrome reveals novel pathways and a strong interaction between the HLA, autoantibodies and the interferon signature. Sci. Rep. 11, 23292. 10.1038/s41598-021-01324-0.

45. Colafrancesco, S., Ciccacci, C., Priori, R., Latini, A., Picarelli, G., Arienzo, F., Novelli, G., Valesini, G., Perricone, C., and Borgiani, P. (2019). STAT4, TRAF3IP2, IL10, and HCP5 Polymorphisms in Sjögren’s Syndrome: Association with Disease Susceptibility and Clinical Aspects. J. Immunol. Res. 2019, 7682827. 10.1155/2019/7682827.

46. Zheng, J., Huang, R., Huang, Q., Deng, F., Chen, Y., Yin, J., Chen, J., Wang, Y., Shi, G., Gao, X., et al. (2015). The GTF2I rs117026326 polymorphism is associated with anti-SSA-positive primary Sjögren’s syndrome. Rheumatol. Oxf. Engl. 54, 562–564. 10.1093/rheumatology/keu466.

47. Casadó-Llombart, S., Gheitasi, H., Ariño, S., Consuegra-Fernández, M., Armiger-Borràs, N., Kostov, B., Ramos-Casals, M., Brito-Zerón, P., and Lozano, F. (2022). Gene Variation at Immunomodulatory and Cell Adhesion Molecules Loci Impacts Primary Sjögren’s Syndrome. Front. Med. 9, 822290. 10.3389/fmed.2022.822290.

48. Emamian, E.S., Leon, J.M., Lessard, C.J., Grandits, M., Baechler, E.C., Gaffney, P.M., Segal, B., Rhodus, N.L., and Moser, K.L. (2009). Peripheral blood gene expression profiling in Sjögren’s syndrome. Genes Immun. 10, 285–296. 10.1038/gene.2009.20.

49. Pranzatelli, T.J.F., Perez, P., Ku, A., Matuck, B., Huynh, K., Sakai, S., Abed, M., Jang, S.-I., Yamada, E., Dominick, K., et al. (2024). GZMK+CD8+ T cells Target A Specific Acinar Cell Type in Sjögren’s Disease. Res. Sq., rs.3.rs-3601404. 10.21203/rs.3.rs-3601404/v2.

50. 50. Urbanski, G., Chabrun, F., Delattre, E., Lacout, C., Davidson, B., Blanchet, O., Chao de la Barca, J.M., Simard, G., Lavigne, C., and Reynier, P. (2023). An immuno-lipidomic signature revealed by metabolomic and machine-learning approaches in labial salivary gland to diagnose primary Sjögren’s syndrome. Front. Immunol. 14, 1205616. 10.3389/fimmu.2023.1205616.

51. Furuzawa-Carballeda, J., Hernández-Molina, G., Lima, G., Rivera-Vicencio, Y., Férez-Blando, K., and Llorente, L. (2013). Peripheral regulatory cells immunophenotyping in primary Sjögren’s syndrome: a cross-sectional study. Arthritis Res. Ther. 15, R68. 10.1186/ar4245.

52. Glauzy, S., Sng, J., Bannock, J.M., Gottenberg, J.-E., Korganow, A.-S., Cacoub, P., Saadoun, D., and Meffre, E. (2017). Defective Early B Cell Tolerance Checkpoints in Sjögren’s Syndrome Patients. Arthritis Rheumatol. Hoboken NJ 69, 2203–2208. 10.1002/art.40215.

53. Xing, Y., Li, B., Wei, P., and Hua, H. (2024). Profiles of peripheral B cell subsets in a cohort of primary Sjögren’s syndrome patients and their potential clinical significance. J. Dent. Sci. 19, 1554–1563. 10.1016/j.jds.2023.12.024.

54. Shiboski, C.H., Baer, A.N., Shiboski, S.C., Lam, M., Challacombe, S., Lanfranchi, H.E., Schiødt, M., Shirlaw, P., Srinivasan, M., Umehara, H., et al. (2018). Natural History and Predictors of Progression to Sjögren’s Syndrome Among Participants of the Sjögren’s International Collaborative Clinical Alliance Registry. Arthritis Care Res. 70, 284–294. 10.1002/acr.23264.

55. Seror, R., Ravaud, P., Bowman, S.J., Baron, G., Tzioufas, A., Theander, E., Gottenberg, J.-E., Bootsma, H., Mariette, X., Vitali, C., et al. (2010). EULAR Sjogren’s syndrome disease activity index: development of a consensus systemic disease activity index for primary Sjogren’s syndrome. Ann. Rheum. Dis. 69, 1103–1109. 10.1136/ard.2009.110619.

56. McGinnis, C.S., Murrow, L.M., and Gartner, Z.J. (2019). DoubletFinder: Doublet Detection in Single-Cell RNA Sequencing Data Using Artificial Nearest Neighbors. Cell Syst. 8, 329–337.e4. 10.1016/j.cels.2019.03.003.

57. Mulè, M.P., Martins, A.J., and Tsang, J.S. (2022). Normalizing and denoising protein expression data from droplet-based single cell profiling. Nat. Commun. 13, 2099. 10.1038/s41467-022-29356-8.

58. Stuart, T., Butler, A., Hoffman, P., Hafemeister, C., Papalexi, E., Mauck, W.M., Hao, Y., Stoeckius, M., Smibert, P., and Satija, R. (2019). Comprehensive Integration of Single-Cell Data. Cell 177, 1888–1902.e21. 10.1016/j.cell.2019.05.031.

59. Borcherding, N., Bormann, N.L., and Kraus, G. (2020). scRepertoire: An R-based toolkit for single-cell immune receptor analysis. F1000Research *9*, 47. 10.12688/f1000research.22139.2.

60. Korsunsky, I., Millard, N., Fan, J., Slowikowski, K., Zhang, F., Wei, K., Baglaenko, Y., Brenner, M., Loh, P., and Raychaudhuri, S. (2019). Fast, sensitive and accurate integration of single-cell data with Harmony. Nat. Methods 16, 1289–1296. 10.1038/s41592-019-0619-0.

61. Karlsson, M., Zhang, C., Méar, L., Zhong, W., Digre, A., Katona, B., Sjöstedt, E., Butler, L., Odeberg, J., Dusart, P., et al. (2021). A single–cell type transcriptomics map of human tissues. Sci. Adv. 7, eabh2169. 10.1126/sciadv.abh2169.

62. Yu, G., Wang, L.-G., Han, Y., and He, Q.-Y. (2012). clusterProfiler: an R Package for Comparing Biological Themes Among Gene Clusters. OMICS J. Integr. Biol. 16, 284–287. 10.1089/omi.2011.0118.

63. Bachmann, M. (2024). rapidfuzz/RapidFuzz: Release 3.8.1 (Zenodo) 10.5281/zenodo.10938887.

64. Jin, S., Guerrero-Juarez, C.F., Zhang, L., Chang, I., Ramos, R., Kuan, C.-H., Myung, P., Plikus, M.V., and Nie, Q. (2021). Inference and analysis of cell-cell communication using CellChat. Nat. Commun. 12, 1088. 10.1038/s41467-021-21246-9.

65. Butler, A., Hoffman, P., Smibert, P., Papalexi, E., and Satija, R. (2018). Integrating single-cell transcriptomic data across different conditions, technologies, and species. Nat. Biotechnol. 36, 411–420. 10.1038/nbt.4096.

