## Supplemental Materials for "Single-cell RNA sequencing of peripheral blood defines two immunological subtypes of Sjögren’s disease"

[1] Department of Orofacial Sciences, University of California, San Francisco, San Francisco, CA, USA
[2] UCSF CoLabs Initiative, University of California, San Francisco, San Francisco, CA, USA
[3] Division of Immunology and Allergology, Laboratory of Translational Immunology, Department of Medicine, Hôpitaux Universitaires de Genève, Geneva, GE, Switzerland
[4] Division of Rheumatology, Department of Medicine, University of California, San Francisco, CA, USA
[5] School of Medicine, University of Washington, Seattle, WA, USA
[6] National Human Genome Research Institute, NIH, Bethesda, MD, USA
[7] Bakar ImmunoX Initiative, University of California, San Francisco, San Francisco, CA, USA
[8] Department of Pathology, University of California, San Francisco, San Francisco, CA, USA
[9] Division of Gastroenterology, Department of Medicine, University of California, San Francisco, San Francisco, CA, USA
[10] Division of Pulmonary, Critical Care, Allergy and Sleep Medicine, University of California, San Francisco, CA, USA
[11] Institute for Human Genetics, University of California, San Francisco, San Francisco, CA, USA
[12] Department of Bioengineering and Therapeutic Sciences, University of California, San Francisco, San Francisco, CA, USA
[13] Department of Epidemiology and Biostatistics, University of California, San Francisco, San Francisco, CA, USA
[14] Chan Zuckerberg Biohub, San Francisco, CA, USA
[15] Parker Institute for Cancer Immunotherapy, San Francisco, CA, USA
[16] Gladstone-UCSF Institute of Genomic Immunology, San Francisco, CA, USA
† These authors contributed equally to this work
‡ These authors contributed equally to this work

<u>Corresponding author</u>: Geoffrey Urbanski. Department of Orofacial Sciences, University of California, San Francisco, San Francisco, CA, USA. UCSF CoLabs Initiative, University of California, San Francisco, San Francisco, CA, USA. Division of Immunology and Allergology, Laboratory of Translational Immunology, Department of Medicine, Hôpitaux Universitaires de Genève, Geneva, GE, Switzerland.

**Supplemental Figure 1. Sample selection and filtering.**

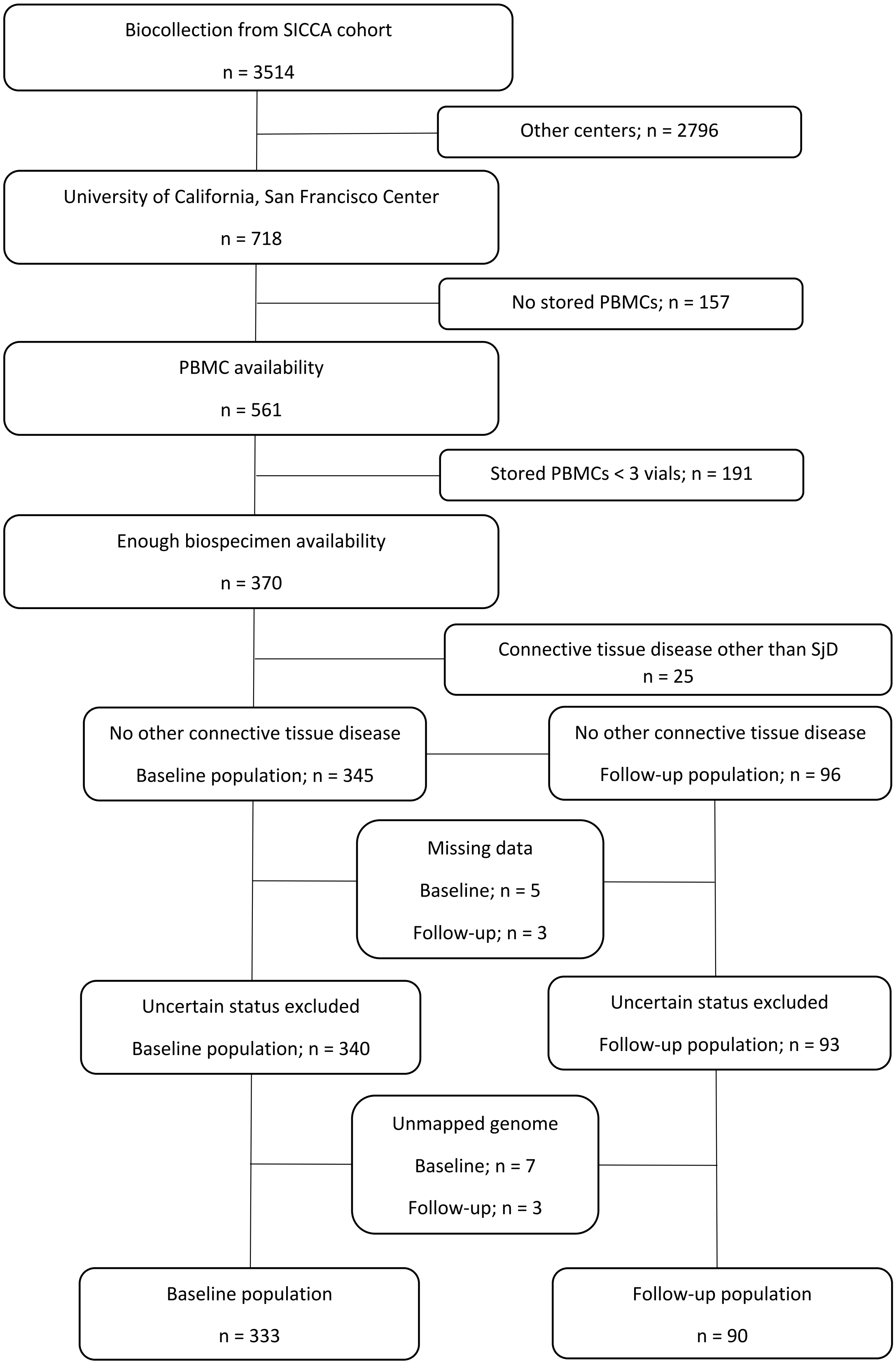

**Supplemental Figure 2. Dotplot of marker genes for annotation of all-cells clusters (Figure 1B Umap).**

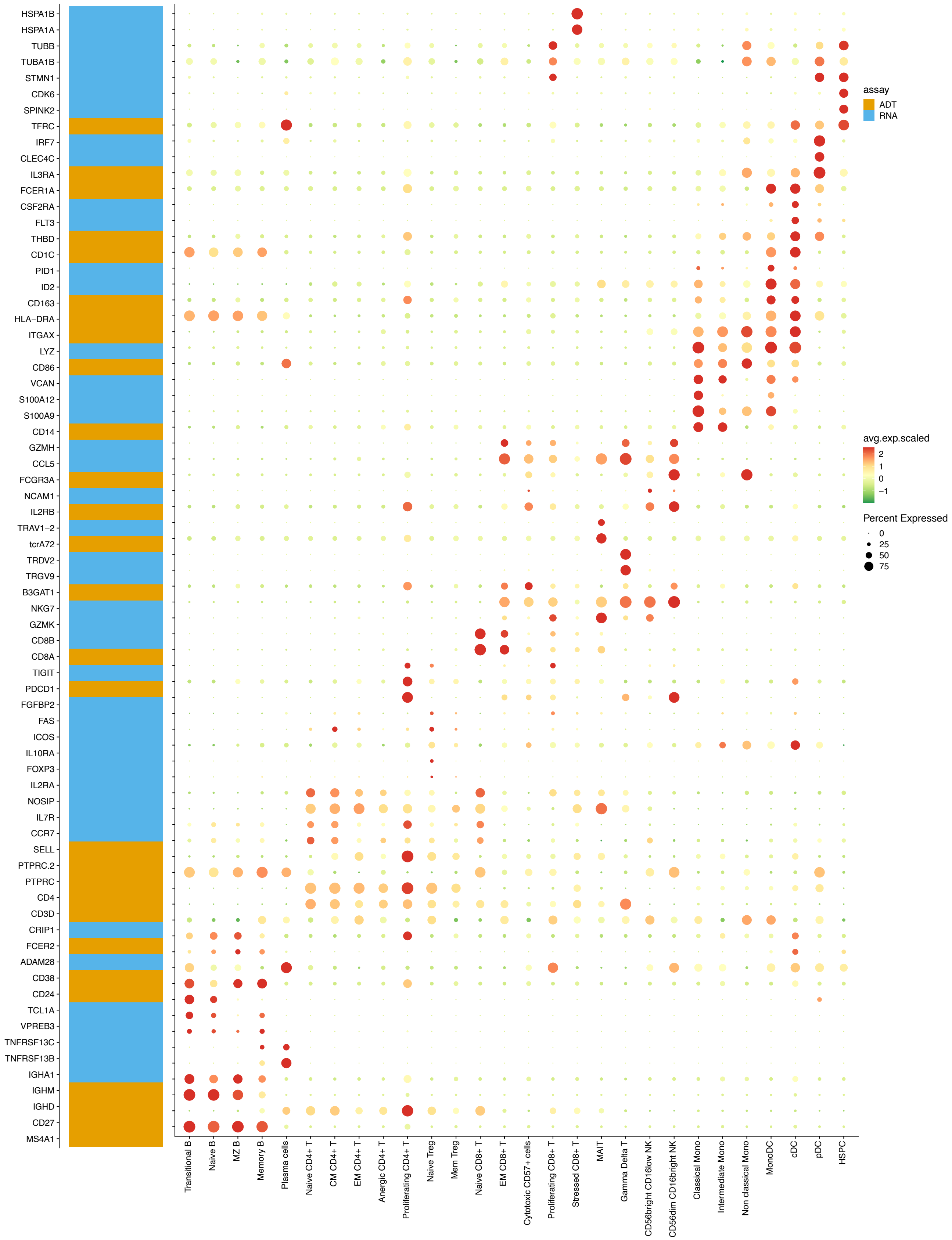

**Supplemental Figure 3. Example volcano plots and comparison of SSA+/SSA- effect sizes for different cell types.** A-C) volcano plots D-F) logFC of SSA+SjD (y-axis) versus SSA-SjD (x-axis); A,D) memory B cells, B,E) classical monocytes, C,F) EM CD4+ T cells.

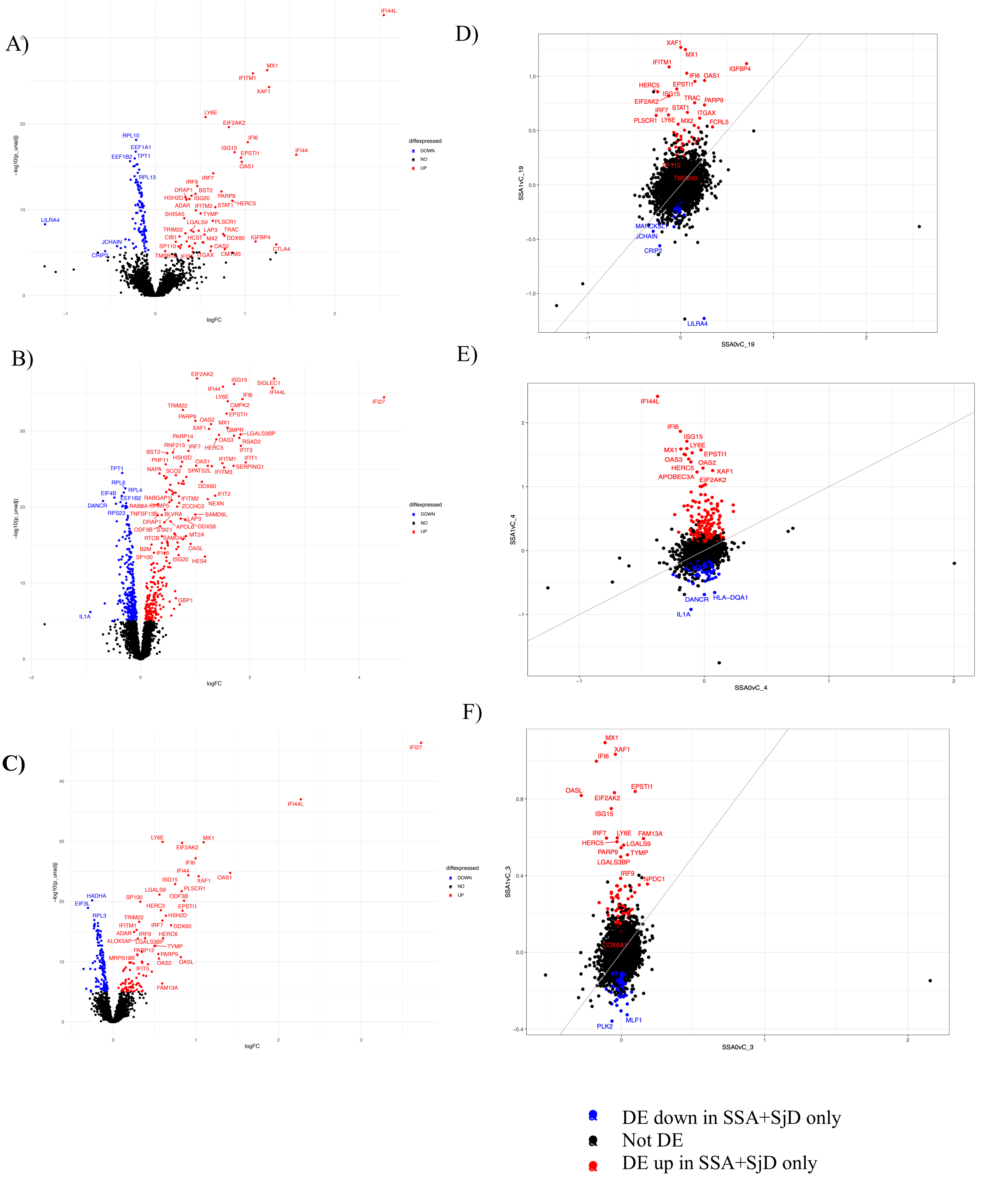

**Supplemental Figure 4. scITD Factor 1 donor score and B cell DE pathways.** A) factor 1 donor score correlation with Chiche IFN modules M1.2, M3.4, and M5.12; B) ACR/EULAR by SSB status, within SSA+ patients; C) scITD factor 1 donor score in SSA+SjD patients with/without immune-modifying medications (including HCQ and Corticosteroids); D) GSEA pathways of B cell "HLA+ISGs" pathway; E) GSEA pathways of B cell "ISGs" pathway.

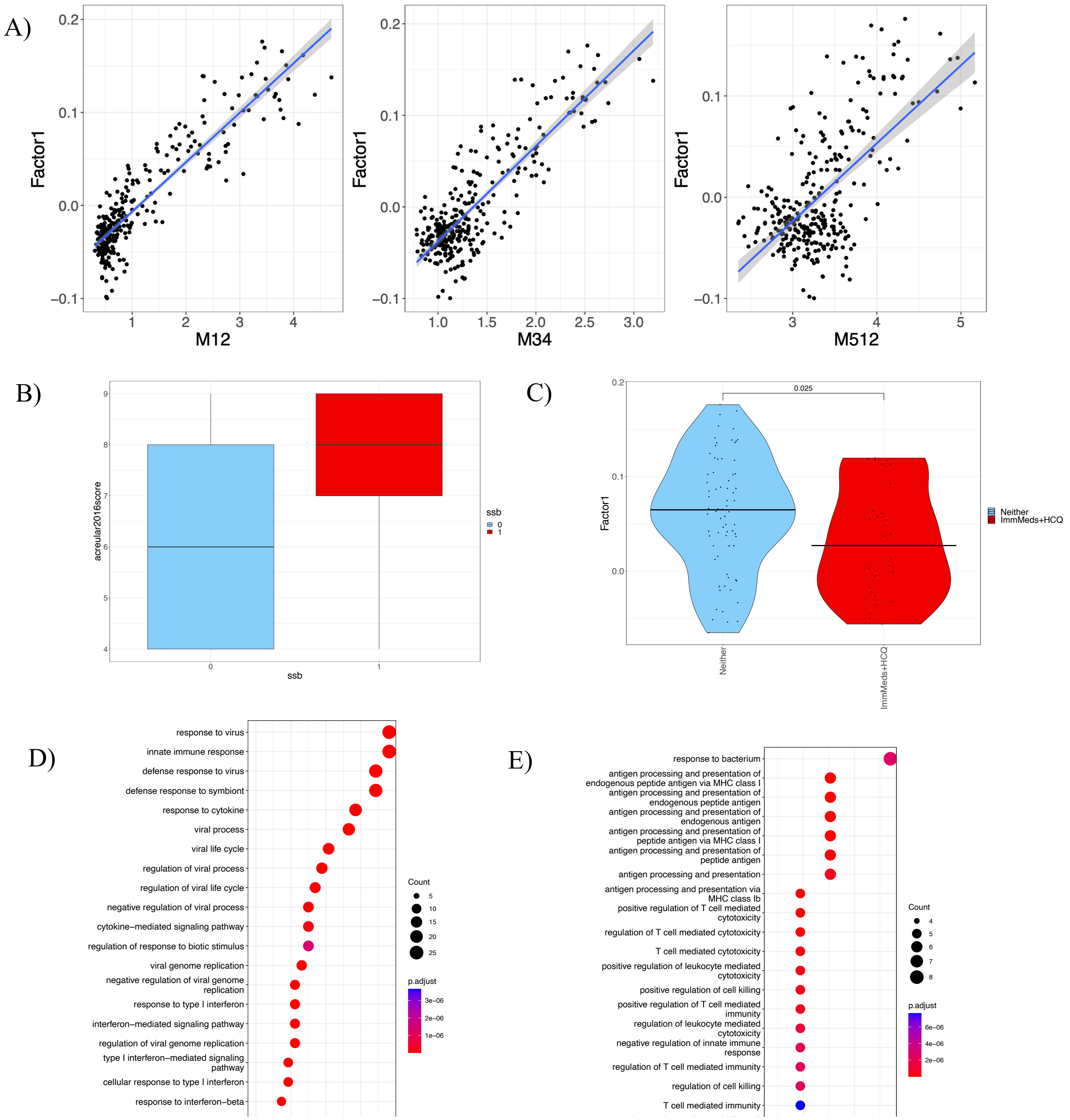

**Supplemental Figure 5. Monocytes and DCs: refined clustering and neighborhood analysis.** A) UMAP of reclustered cells; B) annotation dotplot of marker genes; C) predicted abundances from Dirichlet regression, with significant DA of SSA+SjD and SSA-SjD versus SjD-; D) Milo neighborhood differentials between SSA+SjD and SSA-SjD versus SjD- in B cells; E) Neighborhoods mapped onto the cell clusters with the highest percentage (if >30%) of cells, with logFC of differential abundance, purple=significantly higher than SjD-, pink=significantly lower than SjD-; F) Factor 1 by DA logFC of Milo neighborhoods.

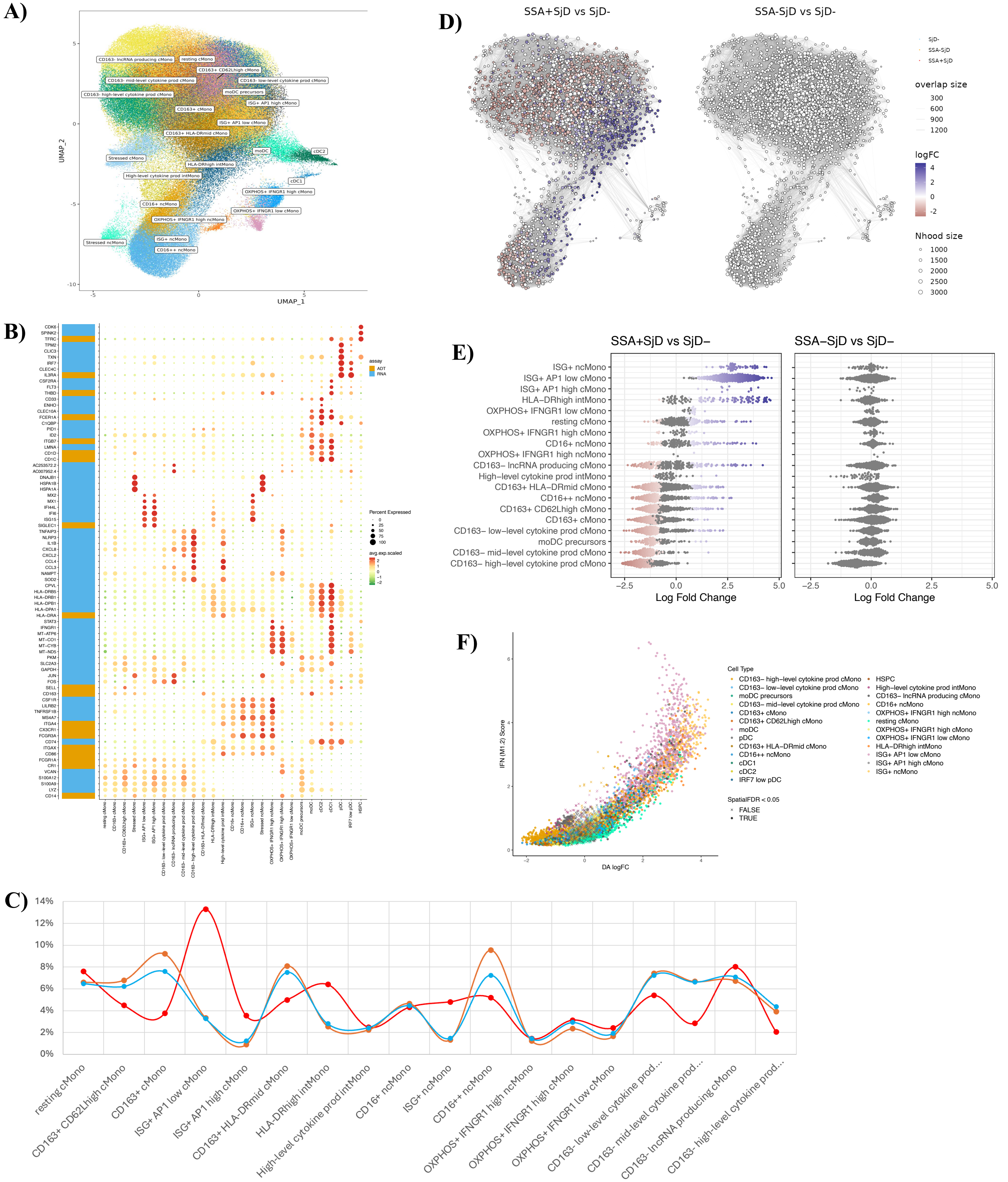

| | resting cMono | CD163+ CD62Lhigh cMono | CD163+ cMono | ISG+ AP1 low cMono | ISG+ AP1 high cMono | CD163+ HLA-DRmid cMono | HLA-DRhigh intMono | High-level cytokine prod intMono | CD16+ ncMono | ISG+ ncMono | CD16++ ncMono | OXPHOS+ IFNGR1 high ncMono | OXPHOS+ IFNGR1 high cMono | OXPHOS+ IFNGR1 low cMono | CD163- low-level cytokine prod... | CD163- mid-level cytokine prod... | CD163- lncRNA producing cMono | CD163- high-level cytokine prod... |
|---|---|---|---|---|---|---|---|---|---|---|---|---|---|---|---|---|---|---|
| **SSA+ sig DA** | | ** | *** | *** | *** | *** | *** | | | *** | ** | | | | ** | *** | | *** |
| **SSA- sig DA** | * | * | ** | | | * | | | | | ** | | | | * | | | |

**Supplemental Figure 6. T & NK cells: Refined clustering and annotation, neighborhood analysis, and TCR length.** A) UMAP of reclustered cells; B) annotation dotplot; C) Milo neighborhood differentials between SSA+SjD and SSA-SjD versus SjD- in B cells; D) Neighborhoods mapped onto the cell clusters with the highest percentage (if >30%) of cells, with logFC of differential abundance; E) Factor 1 by DA logFC of Milo neighborhoods. F) TCR CDR3 beta chain length by SjD/SSA/SSB group.

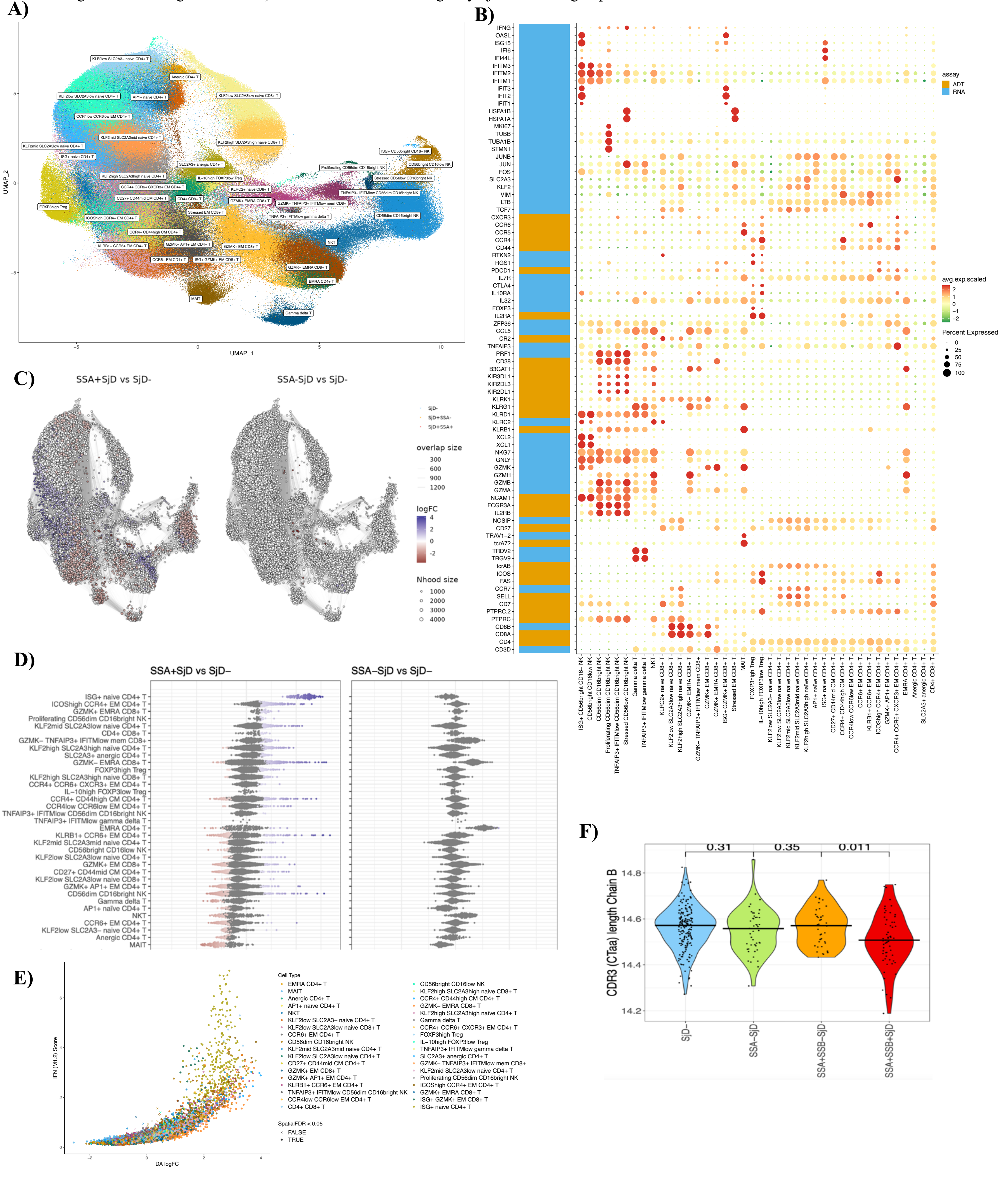

**Supplemental Figure 7: Refined B cells and BCRs.** A-B) IgA and IgG proportion per patient, by SjD/SSA group; C) UMAP with isotypes per cell; D) predicted abundances from Dirichlet regression for lambda cells, by SjD/SSA group; E) percent IGHV3-13 per sample, by group; F) percent IGHV4-34 per sample, by group.

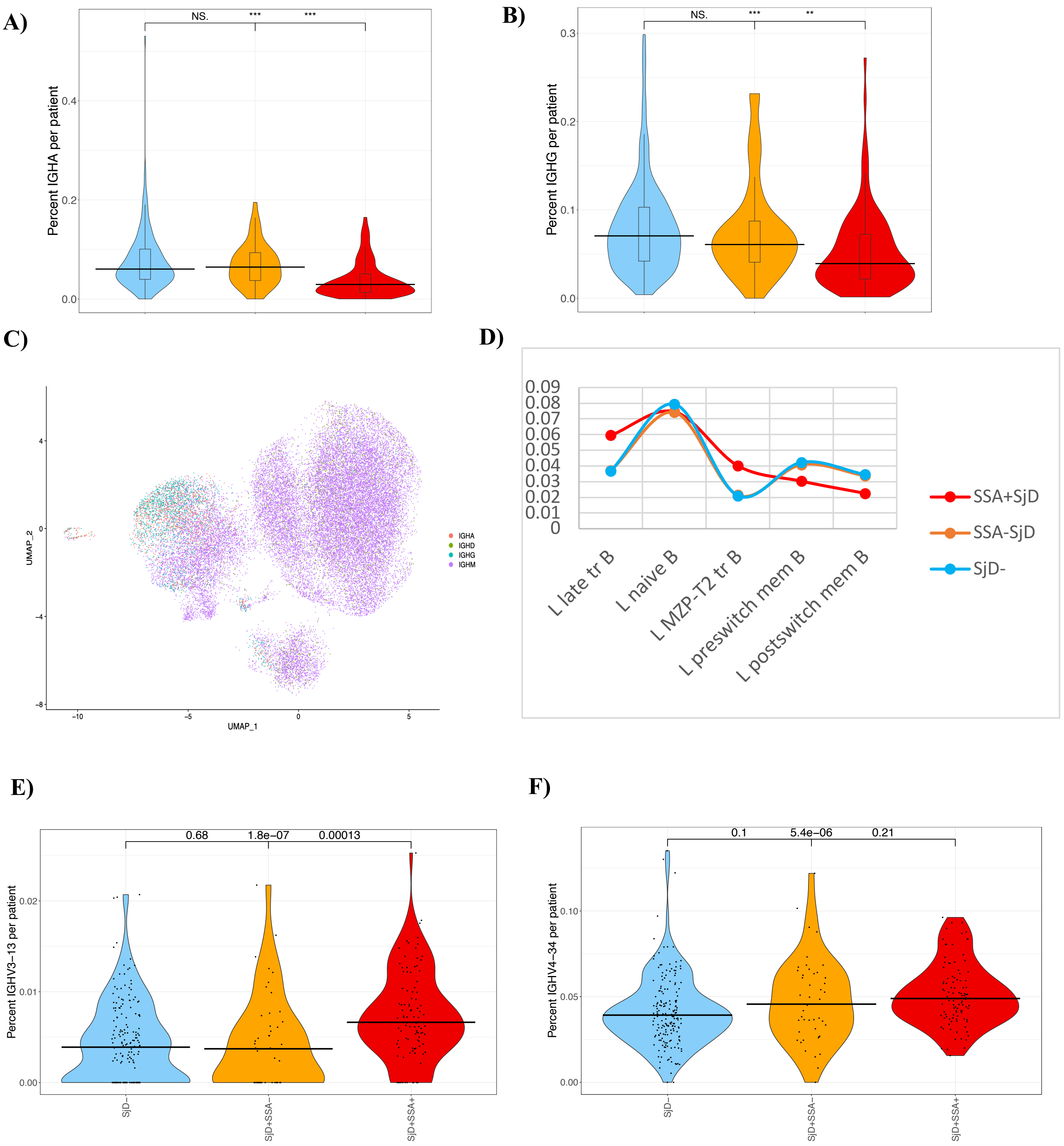

**Supplemental Table 1. Differential abundance via Dirichlet regression and Factor 1 correlation of top-level cell clusters.**

| Cell cluster | SSA+SjD vs SjD- (FDR p) | SSA-SjD vs SjD- (FDR p) | mean SSA+SjD | mean SSA-SjD | mean SjD- | Factor 1 corr | Factor1 (FDR p) |
|---|---|---|---|---|---|---|---|
| **Transitional B** | **4.0E-18** | 0.907 | 0.040 | 0.011 | 0.011 | **0.675** | **4.9E-40** |
| **Naive CD4+ T** | **2.3E-08** | 0.907 | 0.158 | 0.178 | 0.191 | **-0.247** | **0.00029** |
| **Naive CD8+ T** | **3.2E-05** | 0.907 | 0.042 | 0.046 | 0.054 | -0.079 | 1 |
| **EM CD4+ T** | **5.8E-05** | 0.350 | 0.114 | 0.145 | 0.136 | **-0.410** | **3.0E-12** |
| **CM CD4+ T** | **9.8E-05** | 0.907 | 0.044 | 0.024 | 0.028 | **0.570** | **5.0E-26** |
| **Memory B** | **0.000303138** | 0.896 | 0.024 | 0.038 | 0.028 | **-0.273** | **3.0E-05** |
| **MAIT** | **0.00050** | 0.907 | 0.008 | 0.010 | 0.014 | -0.146 | 0.14 |
| **EM CD8+ T** | 0.830 | **0.00061** | 0.096 | 0.111 | 0.095 | -0.019 | 1 |
| **CD56dim CD16bright NK** | **0.00098** | 0.817 | 0.043 | 0.060 | 0.054 | **-0.197** | **0.0096** |
| **MZ B** | **0.023** | 0.907 | 0.013 | 0.008 | 0.007 | **0.316** | **4.4E-07** |
| **Naive B** | **0.028** | 0.896 | 0.068 | 0.048 | 0.048 | **0.212** | **0.0038** |
| **Non classical Mono** | **0.048** | 0.817 | 0.036 | 0.030 | 0.029 | **0.204** | **0.0067** |
| **Proliferating CD8+ T** | 0.070 | 0.907 | 0.0054 | 0.0035 | 0.0029 | **0.438** | **3.3E-14** |
| **pDC** | 0.078 | 0.913 | 0.0028 | 0.0035 | 0.0044 | -0.135 | 0.23 |
| **Naive Treg** | 0.156 | 0.817 | 0.037 | 0.029 | 0.027 | **0.372** | **5.8E-10** |
| **Intermediate Mono** | 0.203 | 0.907 | 0.0088 | 0.0063 | 0.0068 | 0.161 | 0.070 |
| **Gamma Delta T** | 0.238 | 0.907 | 0.0092 | 0.0067 | 0.0093 | -0.067 | 1 |
| **MonoDC** | 0.315 | 0.907 | 0.005 | 0.0058 | 0.006 | **-0.182** | **0.024** |
| **Anergic CD4+ T** | 0.545 | 0.817 | 0.0074 | 0.015 | 0.012 | -0.083 | 1 |
| **CD56bright CD16low NK** | 0.662 | 0.907 | 0.011 | 0.012 | 0.012 | 0.022 | 1 |
| **Plasma cells** | 0.662 | 0.907 | 0.00063 | 0.00025 | 0.00031 | **0.312** | **6.7E-07** |
| **Stressed CD8+ T** | 0.662 | 0.907 | 0.001 | 0.000 | 0.007 | -0.034 | 1 |
| **cDC** | 0.726 | 0.907 | 0.00028 | 0.00039 | 0.00040 | **-0.169** | **0.049** |
| **Classical Mono** | 0.726 | 0.817 | 0.173 | 0.152 | 0.163 | 0.113 | 0.55 |
| **Cytotoxic CD57+ cells** | 0.808 | 0.370 | 0.019 | 0.023 | 0.019 | -0.012 | 1 |
| **HSPC** | 0.808 | 0.907 | 0.00044 | 0.00061 | 0.00049 | -0.110 | 0.57 |
| **Mem Treg** | 0.912 | 0.817 | 0.035 | 0.033 | 0.034 | 0.092 | 1 |
| **Proliferating CD4+ T** | 0.922 | 0.907 | 6.1E-05 | 8.4E-05 | 5.3E-05 | 0.001 | 1 |

**Supplemental Table 2. Interaction and stratified models of Factor 1 with SSA, SSB, and BLA predictors.**

| Patients | Model | Term | Estimate | P-value |
|---|---|---|---|---|
| All | Factor 1 ~ SSA * SSB * BLA | **SSA** | **0.043147** | **9.59E-06** |
| | | SSB | -0.005833 | 0.6454 |
| | | BLA | -0.004720 | 0.4800 |
| | | **SSA*SSB** | **0.041026** | **0.0225** |
| | | **SSA*BLA** | **0.028709** | **0.0166** |
| | | SSB*BLA | -0.013192 | 0.2356 |
| | | SSA*SSB*BLA | NA (collinearity) | NA |
| SSA+ | Factor 1 ~ SSB * BLA | **SSB** | **0.0352** | **0.0450** |
| | | **BLA** | **0.0240** | **0.0795** |
| | | SSB*BLA | -0.0132 | 0.3883 |
| SSA- | Factor 1 ~ SSB * BLA | SSB | -0.0058 | 0.539 |
| | | BLA | -0.0047 | 0.346 |
| | | SSB*BLA | NA (collinearity) | NA |

**Supplemental Table 3. Differential abundances of reclustered Monocytes and Dendritic cells - p-values from Dirichlet regression.**

| | p-fdr Dirichlet vs SjD- | | predicted abundance Dirichlet | | |
|---|---|---|---|---|---|
| Cell cluster | SSA+_fdr | SSA-_fdr | SSA+ | SSA- | SjD- |
| **ISG+ AP1 low cMono** | **6.3E-52** | 0.08 | **13.30%** | 3.33% | **3.25%** |
| **ISG+ ncMono** | **9.2E-27** | 0.45 | **4.80%** | 1.32% | **1.46%** |
| **ISG+ AP1 high cMono** | **8.7E-20** | 0.90 | **3.51%** | 0.87% | **1.23%** |
| **CD163- mid-level cytokine prod cMono** | **8.7E-19** | 0.078 | **2.83%** | 6.65% | **6.61%** |
| **HLA-DRhigh intMono** | **3.3E-15** | 0.45 | **6.40%** | 2.53% | **2.78%** |
| **CD163+ cMono** | **9.6E-15** | **0.00048** | **3.75%** | **9.17%** | **7.57%** |
| **CD163- high-level cytokine prod cMono** | **5.4E-14** | 0.54 | **2.05%** | 3.92% | **4.35%** |
| **moDC precursors** | **8.3E-08** | 0.22 | **2.32%** | 3.76% | **3.96%** |
| **CD163+ HLA-DRmid cMono** | **3.5E-07** | **0.0044** | **4.99%** | **8.07%** | **7.52%** |
| **CD16++ ncMono** | **2.1E-05** | **2.2E-05** | **5.18%** | **9.53%** | **7.25%** |
| **CD163+ CD62Lhigh cMono** | **5.0E-05** | **0.0015** | **4.46%** | **6.77%** | **6.22%** |
| **CD163- low-level cytokine prod cMono** | **0.00014** | **0.0092** | **5.38%** | **7.37%** | **7.22%** |
| pDC | **0.008** | 0.83 | **2.29%** | 2.37% | **3.06%** |
| moDC | **0.024** | 0.45 | **2.76%** | 3.09% | **3.40%** |
| OXPHOS+ IFNGR1 low cMono | 0.10 | 0.45 | 2.41% | 1.63% | 1.91% |
| cDC2 | 0.14 | 0.66 | 1.91% | 1.84% | 2.16% |
| IRF7 low pDC | 0.22 | 0.90 | 1.09% | 0.94% | 1.30% |
| resting cMono | 0.31 | **0.030** | 7.59% | **6.58%** | 6.48% |
| cDC1 | 0.47 | 0.66 | 0.94% | 0.85% | 1.04% |
| CD16+ ncMono | 0.53 | 0.075 | 4.31% | 4.64% | 4.45% |
| HSPC | 0.61 | 0.73 | 1.11% | 0.96% | 1.18% |
| CD163- lncRNA producing cMono | 0.69 | 0.11 | 7.99% | 6.71% | 7.10% |
| OXPHOS+ IFNGR1 high cMono | 0.70 | 0.73 | 3.12% | 2.33% | 2.91% |
| High-level cytokine prod intMono | 0.70 | 0.48 | 2.47% | 2.23% | 2.43% |
| OXPHOS+ IFNGR1 high ncMono | 0.86 | 0.66 | 1.43% | 1.24% | 1.43% |

**Supplemental Table 4. Differential abundances of reclustered T and NK cells - p-values from Dirichlet regression.**

| | SSA+_fdr | SSA-_fdr | mean SSA+SjD | mean SSA-SjD | mean SjD- |
|---|---|---|---|---|---|
| **ISG+ naive CD4+ T** | **1.82E-27** | 0.956 | **2.13%** | 0.60% | **0.66%** |
| **CCR6+ EM CD4+ T** | **2.04E-09** | 0.956 | **2.75%** | 4.18% | **4.25%** |
| **AP1+ naive CD4+ T** | **2.31E-07** | 0.764 | **1.68%** | 2.12% | **2.65%** |
| **KLF2mid SLC2A3mid naive CD4+ T** | **3.94E-07** | 0.887 | **4.45%** | 5.33% | **6.14%** |
| **GZMK- EMRA CD8+ T** | **5.03E-07** | **0.001** | **4.39%** | 3.80% | **2.60%** |
| **KLF2low SLC2A3low naive CD4+ T** | **1.90E-06** | 0.956 | **5.58%** | 6.98% | **7.20%** |
| **ICOShigh CCR4+ EM CD4+ T** | **2.72E-06** | 0.887 | **3.49%** | 2.20% | **2.12%** |
| **KLF2low SLC2A3- naive CD4+ T** | **2.72E-06** | 0.590 | **2.17%** | 3.56% | **3.22%** |
| **KLF2low SLC2A3low naive CD8+ T** | **1.99E-05** | 0.887 | **2.86%** | 3.60% | **4.10%** |
| **MAIT** | **3.78E-05** | 0.887 | **1.08%** | 1.25% | **1.63%** |
| **CD27+ CD44mid CM CD4+ T** | **4.42E-05** | 0.887 | **3.49%** | 4.56% | **4.47%** |
| **GZMK+ EMRA CD8+ T** | **0.003** | 0.956 | **1.16%** | 0.74% | **0.75%** |
| **EMRA CD4+ T** | 0.953 | **0.034** | 1.15% | **1.45%** | **1.02%** |
| **CCR4low CCR6low EM CD4+ T** | 0.638 | **0.036** | 3.18% | **3.53%** | **2.87%** |
| **FOXP3high Treg** | **0.046** | 0.887 | **4.20%** | 3.39% | **3.24%** |
| Proliferating CD56dim CD16bright NK | 0.075 | 0.956 | 0.77% | 0.59% | 0.57% |
| SLC2A3+ anergic CD4+ T | 0.078 | 0.887 | 3.37% | 2.75% | 2.76% |
| CD56dim CD16bright NK | 0.080 | 0.887 | 6.37% | 7.23% | 6.75% |
| KLF2mid SLC2A3low naive CD4+ T | 0.149 | 0.956 | 5.65% | 4.47% | 4.60% |
| NKT | 0.149 | 0.764 | 2.03% | 2.26% | 2.07% |
| GZMK- TNFAIP3+ IFITMlow mem CD8+ | 0.158 | 0.641 | 2.05% | 1.87% | 1.61% |
| GZMK+ EM CD8+ T | 0.293 | 0.764 | 5.22% | 5.60% | 5.31% |
| Gamma delta T | 0.480 | 0.956 | 1.14% | 0.98% | 1.24% |
| GZMK+ AP1+ EM CD4+ T | 0.489 | 0.956 | 4.52% | 4.42% | 4.48% |
| KLF2high SLC2A3high naive CD8+ T | 0.489 | 0.956 | 3.43% | 2.84% | 3.66% |
| KLRC2+ naive CD8+ T | 0.489 | 0.956 | 0.44% | 0.42% | 0.47% |
| Anergic CD4+ T | 0.535 | 0.887 | 0.63% | 0.73% | 0.68% |
| CCR4+ CD44high CM CD4+ T | 0.638 | 0.956 | 4.20% | 3.79% | 3.74% |
| KLF2high SLC2A3high na√Øve CD4+ T | 0.638 | 0.764 | 4.06% | 3.03% | 3.63% |
| CD4+ CD8+ T | 0.640 | 0.969 | 0.90% | 0.73% | 0.79% |
| KLRB1+ CCR6+ EM CD4+ T | 0.653 | 0.764 | 2.97% | 2.94% | 2.66% |
| TNFAIP3+ IFITMlow CD56dim CD16bright NK | 0.653 | 0.887 | 1.51% | 1.66% | 1.52% |
| Stressed CD56low CD16bright NK | 0.663 | 0.956 | 0.28% | 0.27% | 0.29% |
| CCR4+ CCR6+ CXCR3+ EM CD4+ T | 0.712 | 0.956 | 2.13% | 1.96% | 1.98% |
| Stressed EM CD8+ T | 0.713 | 0.956 | 0.34% | 0.30% | 0.34% |
| ISG+ GZMK+ EM CD8+ T | 0.721 | 0.997 | 0.44% | 0.37% | 0.38% |
| TNFAIP3+ IFITMlow gamma delta T | 0.721 | 0.956 | 0.46% | 0.38% | 0.42% |
| ISG+ CD56bright CD16- NK | 0.761 | 0.956 | 0.32% | 0.29% | 0.29% |
| CD56bright CD16low NK | 0.903 | 0.956 | 1.97% | 1.81% | 1.85% |
| IL-10high FOXP3low Treg | 0.905 | 0.956 | 1.04% | 1.01% | 0.98% |

**Supplemental Table 5. Differential abundances of reclustered B cells from Dirichlet regression.**

| | p_fdr vs SjD- | | predicted abundance from Dirichlet regression | | |
|---|---|---|---|---|---|
| | SSA+ | SSA- | SSA+SjD | SSA-SjD | SjD- |
| **Resting K mem B** | **6.5E-19** | **0.027** | **2.39%** | **5.16%** | **5.74%** |
| **Activated ISG+ K tr B** | **1.3E-13** | 0.986 | 3.89% | 2.13% | 1.68% |
| **K postswitch mem B** | **1.2E-12** | **0.029** | **3.63%** | **6.21%** | **6.73%** |
| **K preswitch mem B** | **2.9E-12** | **0.040** | **3.16%** | **5.56%** | **5.99%** |
| **K MZP-T2 tr B** | **3.2E-10** | 0.986 | 4.48% | 2.91% | 2.31% |
| **L MZP-T2 tr B** | **6.9E-09** | 0.240 | 4.01% | 2.14% | 2.11% |
| **ISG+ Kappa naive B** | **9.1E-07** | 0.821 | 1.21% | 0.78% | 0.64% |
| **K naive B** | **1.4E-06** | **0.00033** | **12.98%** | **14.88%** | **16.53%** |
| **L postswitch mem B** | **9.4E-06** | 0.144 | 2.26% | 3.37% | 3.46% |
| **L late tr B** | **1.9E-05** | 0.158 | 5.95% | 3.70% | 3.69% |
| **K late tr B** | **6.2E-05** | 0.077 | 8.05% | 5.25% | 5.43% |
| **L preswitch mem B** | **0.00031** | 0.101 | 3.03% | 4.09% | 4.23% |
| **K early naive B** | **0.00035** | 0.139 | 14.71% | 11.47% | 10.58% |
| **K DN B** | **0.00035** | **0.024** | **3.33%** | **4.13%** | **4.52%** |
| **CD27+ ADAM28+ AHNAK+ MZ B** | **0.0017** | 0.158 | 1.47% | 2.02% | 2.08% |
| **CD27- FCER2high ADAM28+ MZP** | **0.0029** | 0.986 | 4.28% | 3.95% | 3.10% |
| **L naive B** | 0.084 | **0.018** | **7.46%** | **7.42%** | **7.96%** |
| CD11c+ T-bet+ mem B | 0.034 | 0.986 | 3.65% | 5.56% | 4.32% |
| TXNIP+ K naive B | 0.10 | 0.198 | 3.34% | 2.81% | 2.75% |
| CD27- FCER2mid ADAM28+ MZP | 0.40 | 0.153 | 3.97% | 3.74% | 3.57% |
| FAS+ CD11c+ T-bet+ mem B | 0.72 | 0.156 | 1.71% | 1.68% | 1.70% |
| Plasma cells | 0.34 | 0.821 | 1.03% | 1.05% | 0.88% |

**Supplemental Table 6. Mean isotype proportions by SjD/SSA group.**

| | **SjD-** | **SjD+SSA-** | **SjD+SSA+** | **p SSA+ vs SjD-** |
|---|---|---|---|---|
| **IgA*** | 0.0749 | 0.0717 | **0.0372** | **2.90E-13** |
| **IgD** | 0.0647 | 0.0664 | 0.0609 | 0.47 |
| **IgG*** | 0.0775 | 0.0770 | **0.0504** | **9.10E-08** |
| **IgM** | 0.784 | 0.785 | **0.851** | **7.60E-11** |

**Supplemental Table 7. Differences in heavy chain CDR3 length by SjD/SSA group. (t-test)**

| | SSA+Pfdr | SSA-P | meanSSA+ | meanSSA- | meanSjD- |
|---|---|---|---|---|---|
| **K naive B** | **7.9E-08** | 0.626 | **17.809** | 18.614 | **18.532** |
| **K early naive B** | **1.2E-06** | 0.271 | **17.860** | 18.404 | **18.596** |
| **TXNIP+ K naive B** | **0.0026** | 0.324 | **17.983** | 18.568 | **19.081** |
| K late tr B | 0.3320 | 0.120 | 18.112 | 18.944 | 18.489 |
| L naive B | 0.3629 | 0.787 | 17.994 | 18.430 | 18.356 |
| L MZP-T2 tr B | 1 | 0.563 | 18.352 | 18.531 | 18.827 |
| L late tr B | 1 | 0.934 | 18.171 | 18.546 | 18.511 |
| CD27- FCER2mid ADAM28+ MZP | 1 | 0.701 | 17.741 | 18.412 | 18.283 |
| CD27- FCER2high ADAM28+ MZP | 1 | 0.039 | 17.773 | 19.032 | 18.141 |
| K DN B | 1 | 0.229 | 17.879 | 18.472 | 18.131 |
| Activated ISG+ K tr B | 1 | 0.778 | 18.138 | 18.301 | 18.455 |
| CD11c+ T-bet+ mem B | 1 | 0.290 | 17.609 | 17.718 | 17.364 |
| ISG+ Kappa naive B | 1 | 0.365 | 18.319 | 17.750 | 18.904 |
| Plasma cells | 1 | 0.601 | 17.221 | 17.174 | 16.835 |
| FAS+ CD11c+ T-bet+ mem B | 1 | 0.012 | 17.970 | 16.590 | 17.740 |
| L postswitch mem B | 1 | 0.497 | 17.445 | 17.407 | 17.582 |
| K MZP-T2 tr B | 1 | 0.407 | 18.493 | 18.972 | 18.631 |
| Resting K mem B | 1 | 0.437 | 17.064 | 16.805 | 16.973 |
| L preswitch mem B | 1 | 0.202 | 17.153 | 16.904 | 17.224 |
| CD27+ ADAM28+ AHNAK+ MZ B | 1 | 0.165 | 17.223 | 16.637 | 17.286 |
| K postswitch mem B | 1 | 0.439 | 17.398 | 17.581 | 17.419 |
| K preswitch mem B | 1 | 0.948 | 16.863 | 16.884 | 16.869 |

**Supplemental Table 8. Cell types with significant differences in IGHV4-34 and IGHV3-13 usage.**

| | SSA+SjD vs SjD- (FDR p) | SSA-SjD vs SjD- (FDR p) | mean SSA+SjD | mean SSA-SjD | mean SjD- |
|---|---|---|---|---|---|
| **IGHV4-34** | | | | | |
| L MZP-T2 tr B | 0.00089 | 1 | 0.054 | 0.043 | 0.032 |
| K naive B | 0.0063 | 1 | 0.056 | 0.056 | 0.041 |
| K MZP-T2 tr B | 0.014 | 1 | 0.080 | 0.064 | 0.059 |
| L late tr B | 0.015 | 1 | 0.071 | 0.067 | 0.045 |
| Activated ISG+ K tr B | 0.020 | 1 | 0.073 | 0.072 | 0.065 |
| **IGHV3-13** | | | | | |
| K early naive B | 6.9E-07 | 1 | 0.0076 | 0.0072 | 0.0026 |
| Activated ISG+ K tr B | 7.4E-05 | 1 | 0.0119 | 0.0028 | 0.0056 |
| L naive B | 0.0012 | 1 | 0.0101 | 0.0054 | 0.0032 |
| K MZP-T2 tr B | 0.0015 | 1 | 0.0045 | 0.0000 | 0.0020 |
| K late tr B | 0.0025 | 1 | 0.0089 | 0.0012 | 0.0062 |
| L MZP-T2 tr B | 0.023 | 1 | 0.0064 | 0.0017 | 0.0069 |
| CD27- FCER2high ADAM28+ MZP | 0.023 | 1 | 0.0083 | 0.0021 | 0.0046 |

**Supplemental Table 9. Cell type marker genes.**

| Cell type | Abbreviations | Markers |
|---|---|---|
| Transitional B | Tr B | CD19, CD20, CD27, IGHD, IGHM, TNFRSF13C, VPREB3, TCL1A, CD24, CD38 |
| Marginal Zone Precursors / Type 2 Transitional B | MZP-T2 | CD19, CD20, CD27, IGHD, IGHM, TNFRSF13C, VPREB3, TCL1A, CD24, CD38, MZB1, PLD4 |
| Marginal Zone Precursors | MZP | CD19, CD20, CD27, IGHD, IGHM, TNFRSF13C, ADAM28, FCER2 |
| Naïve B | | CD19, CD20, CD27, IGHD, IGHM, TNFRSF13C, TCL1A |
| Marginal Zone (MZ) B | MZ B | CD19, CD20, CD27, IGHD, IGHM, TNFRSF13C, ADAM28, FCER2, AHNAK |
| Memory B | Mem B | CD19, CD20, CD27, IGHM, IGHA1, TNFRSF13B, TNFRSF13C, CRIP1 |
| Plasma cells | | CD19, CD20, CD27, TNFRSF13B, CRIP1, CD38 |
| Naive CD4+ T | | CD3D, CD4, CD45RA (PTPRC), CD27, SELL, CCR7, IL7R, NOSIP |
| Central Memory CD4+T | CM CD4+ T | CD3D, CD4, CD45RO (PTPRC.2), CD27, SELL, CCR7, IL7R, NOSIP, ICOS, FAS |
| Effector Memory CD4+ T | EM CD4+ T | CD3D, CD4, CD45RO (PTPRC.2), CD27, SELL, CCR7, IL7R, NOSIP, ICOS, FAS |
| Regulatory CD4+ T | Treg | CD3D, CD4, CD45RO (PTPRC.2), CD27, IL2RA, FOXP3, IL10RA, ICOS, FAS, PDCD1 |
| Naive CD8+ T | | CD3D, CD8A, CD8B, CD45RA (PTPRC), CD27, SELL, CCR7, IL7R, NOSIP |
| Effector Memory CD8+ T | EM CD8+ T | CD3D, CD8A, CD8B, CD45RO (PTPRC.2), CD27, SELL, CCR7, GZMH, GZMK, NKG7 |
| Effector Memory CD8+ T expressing CD45RA | EMRA CD8+ T | CD3D, CD8A, CD8B, CD45RA (PTPRC), CD27, SELL, CCR7, GZMH, GZMK, NKG7 |
| Mucosal-associated invariant T | MAIT | TCRA7.2, TRAV1-2, IL7R |
| Gamma Delta T | | CD3D, TRGV9, TRDV2, CCL5 |
| CD56 bright CD16 low NK | | IL2RB, NCAM1, FCGR3A, NKG7, CCL5, GZMH |
| CD56 dim CD16 bright NK | | IL2RB, NCAM1, FCGR3A, NKG7, CCL5, GZMH |
| Classical monocytes | cMono | CD14, S100A9, S100A12, VCAN, LYZ |
| Intermediate monocytes | intMono | CD14, FCGR3A, S100A9, VCAN, LYZ, HLA-DRA, CD86, ITGAX |

| | | |
|---|---|---|
| Non-classical monocytes | ncMono | CD14, FCGR3A, S100A9, LYZ, HLA-DRA, CD86, ITGAX |
| Monocyte-derived DCs | MoDC | CD14, S100A9, S100A12, VCAN, LYZ HLA-DRA, CD163, ID2, PID1 |
| Conventional dendritic cells | cDC | CD1C, THBD, FLT3, CSF2RA, FCER1A, HLA-DRA |
| Type1 conventional dendritic cells | cDC1 | CD1C, THBD, FLT3, CSF2RA, FCER1A, HLA-DRA, HLA-DPA1, HLA-DRB1, CXCR1, ITGA4 |
| Type2 conventional dendritic cells | cDC2 | CD1C, C1QBP, ENHO, CD33 |
| Plasmacytoid dendritic cells | pDC | IL3RA, CLEC4C, IRF7 |
| Hematopoietic stem and progenitor cells | HSPC | TFRC, SPINK2, CDK6 |
| **Specific state** | **Abbreviations** | **Markers** |
| Activated | AP1 | Both FOS and JUN |
| Anergic/Resting | | No upregulated gene |
| Exhausted | | PDCD1, FGFBP2, TIGIT |
| Kappa light chain | K | IGKC |
| Lambda light chain | L | IGLC2, IGLC3 |
| Preswitched | preswitch | IGHM high, IGHA1 low |
| Postswitched | postswitch | IGHM low or null, IGHA1 high |
| Proliferating | | TUBA1B, TUBB, STMN1 |
| Stressed | | HSP1A, HSP1B |
| Upregulation of Interferon-Stimulated Genes | ISG+ | Combination within IFIT1, IFIT2, IFIT3, IFI44L, IFI6, ISG15, OASL |
